# Fast *Klebsiella pneumoniae* typing for outbreak reconstruction: an highly discriminatory HRM protocol on *wzi* capsular gene developed using EasyPrimer tool

**DOI:** 10.1101/679001

**Authors:** Matteo Perini, Aurora Piazza, Simona Panelli, Domenico Di Carlo, Marta Corbella, Floriana Gona, Francesca Vailati, Piero Marone, Daniela Maria Cirillo, Claudio Farina, Gian Vincenzo Zuccotti, Francesco Comandatore

## Abstract

In this work we present EasyPrimer, a user-friendly online tool developed to assist pan-PCR and High Resolution Melting (HRM) primer design. The tool finds the most suitable regions for primer design in a gene alignment and returns a clear graphical representation of their positions on the gene. EasyPrimer is particularly useful in difficult contexts, e.g. on gene alignments of hundreds of sequences and/or on highly variable genes. HRM analysis is an emerging method for fast and cost saving bacterial typing and an HRM scheme of six primer sets on five Multi-Locus Sequence Type (MLST) genes is already available for *Klebsiella pneumoniae*. We validated the tool designing a scheme of two HRM primer sets on the hypervariable gene *wzi* of *Klebsiella pneumoniae* and compared the two schemes. The *wzi* scheme resulted to have a discriminatory power comparable to the HRM MLST scheme, using only one third of primer sets. Then we successfully used the *wzi* HRM primer scheme to reconstruct a *Klebsiella pneumoniae* nosocomial outbreak in few hours. The use of hypervariable genes reduces the number of HRM primer sets required for bacterial typing allowing to perform cost saving, large-scale surveillance programs.

## Introduction

Most methods used for the identification and typing of prokaryotes are based on DNA amplification and sequencing. Indeed, the sequence of specific genes can harbour enough information to classify bacteria at both species and subspecies level. For instance, Multi-Locus Sequence Typing (MLST) is one of the most used methods for bacterial typing and it is based on the amplification and sequencing of few housekeeping genes^1^. During the last ten years, the analysis of the entire bacterial genome by Whole Genome Sequencing (WGS) approach revolutionized the field, drastically increasing the typing precision^1^.

The reconstruction of nosocomial outbreaks is one of the most important clinical applications of bacterial typing. A nosocomial outbreak occurs when the number of patients infected by a pathogen increases above the expected in a limited time^2^. In these situations, it is fundamental to determine the clonality of bacteria causing disease in the patients to define the proper strategy to handle the emergency. Pulsed-Field Gel Electrophoresis (PFGE), MLST and WGS are the most frequently applied molecular methods in outbreak investigation^1^.

During a nosocomial outbreak, clinicians need bacterial typing information in the shortest time possible. Despite the high potential of WGS in outbreak reconstruction, the sequencing of a complete genome requires two to four working days, introducing an important time lag. Similarly, PFGE typing requires five days and also MLST needs few days. During the last years, the High Resolution Melting (HRM) assay has emerged as a low-cost and fast method for bacterial typing, particularly promising for epidemiological applications^3,4,5,6^. HRM is a single-step procedure for the discrimination of sequence variants on the basis of their melting temperature. This method allows to perform bacterial typing in less than five hours^7^. To develop a novel HRM-based typing procedure, it is necessary to: i) select one or more core genes; ii) design a primer set in conserved regions flanking a gene portion where the melting temperature varies among the strains.

Andersson and colleagues have developed the “MinimumSNP” tool^8^, which identifies, in a gene alignment, the variable positions that can lead to a melting temperature change (called informative SNPs). MinimumSNP identifies single informative positions, that could be spread along the entire alignment. In other words, it does not indicate which regions are more suitable for primer design: two low-variable regions neighbouring a SNP-rich, informative stretch. Thus, the user has to choose one (or few) SNPs and then design primers around it (or around them).

Herein, we present EasyPrimer, a web-based tool for the identification of the gene regions suitable for primers design to perform HRM studies and any kind of pan-PCR experiments. Moreover, we validated EasyPrimer by designing HRM primers for the discrimination of clinical isolates of *Klebsiella pneumoniae*, an important opportunistic pathogen frequently cause of infections in humans and animals^9^.

## Results

### EasyPrimer: a tool for primers design

EasyPrimer is a user-friendly open-source tool developed to assists primer design in difficult contexts, e.g. on an alignment of hundreds of sequences and/or on hypervariable genes. The tool uses as input a sequence alignment and identifies the best regions for primer design: two low variable regions flanking a highly variable one. The on-line and the stand-alone versions of the tool are freely available at https://skynet.unimi.it/index.php/tools/easyprimer.

### Primers design

We downloaded *pgi*, *gapA* and *wzi* gene sequences from BigsDB database (https://bigsdb.pasteur.fr) and we run EasyPrimer to identify the best regions for primer design. The EasyPrimer output for the *wzi* gene is reported in Figure 1, while the outputs relative to *pgi* and *gapA* genes are reported in <u>Supplementary Fig. S1</u> and <u>Supplementary Fig. S2</u>, respectively. Then we designed a total of four primer sets: one for *pgi*, one for *gapA* and two for *wzi* (reported in Table 1).

**Table 1.**
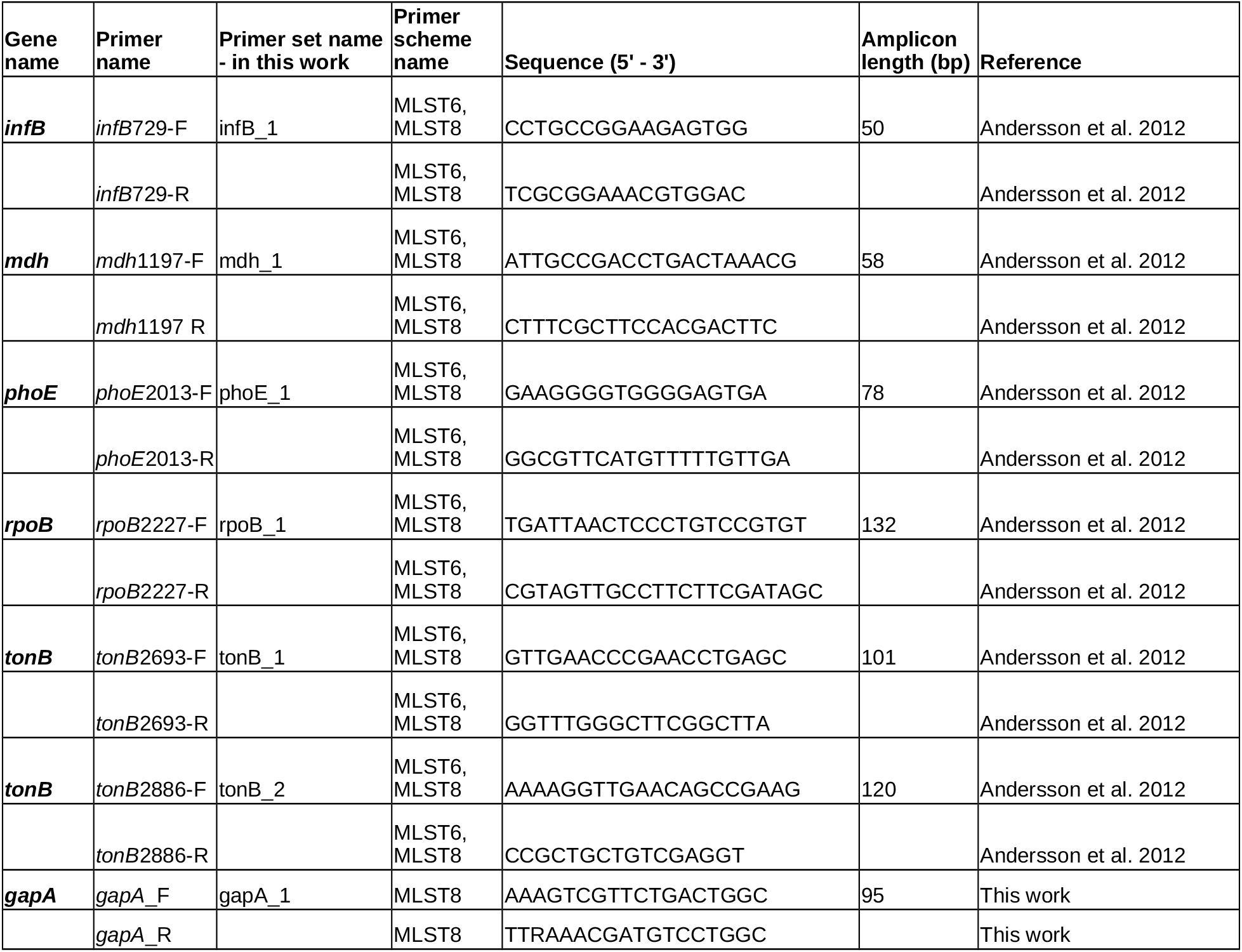
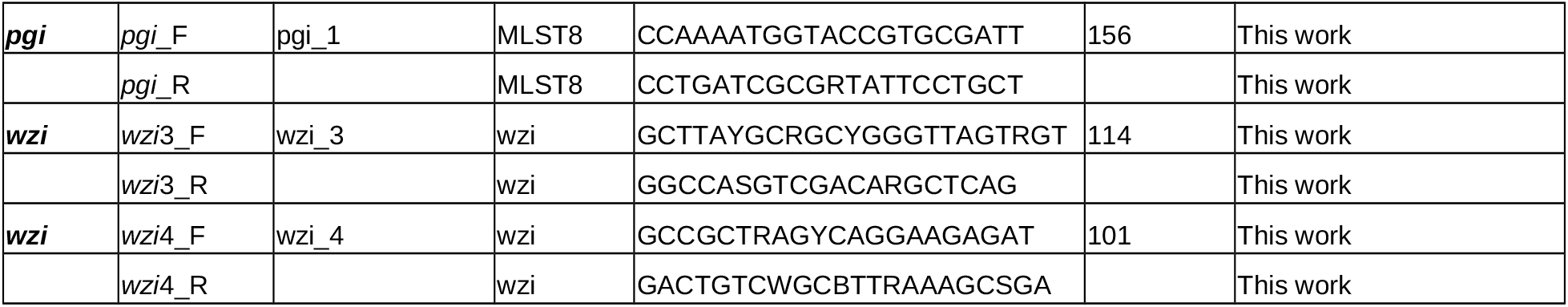
Primer sets used in this work.

**Figure 1.**
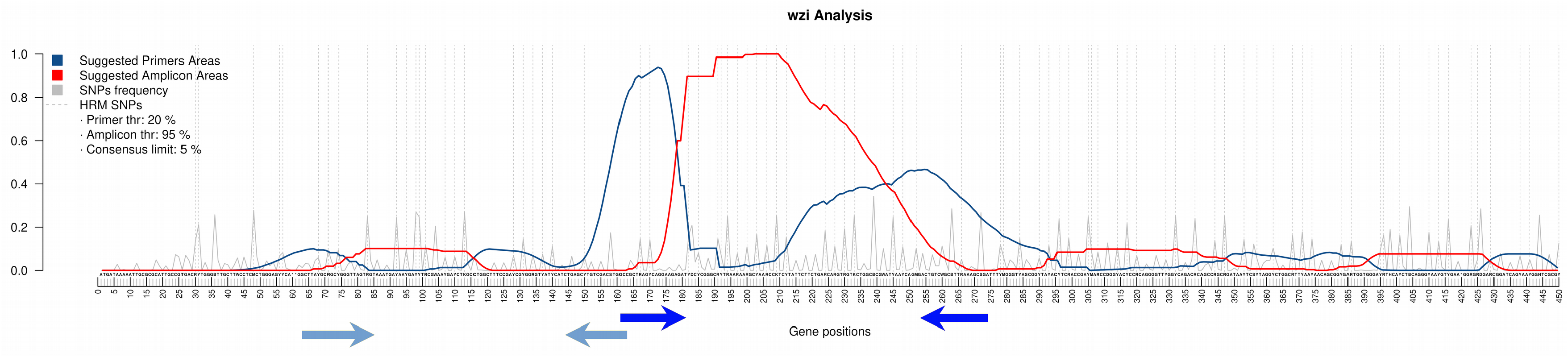
EasyPrimer output for the *wzi gene* (563 alleles on https://bigsdb.pasteur.fr). The consensus sequence calculated from the gene alignment is reported on the x-axis. Residues under the peaks of the blue curve are highly conserved and thus suitable for primer design. Conversely the red curve increases over the highly variable regions suggested to be amplified. The grey peaks represent all the Single Nucleotide Polymorphisms (SNPs) with their own frequency. The dotted lines are used to highlight the “HRM-detectable” SNPs, i.e. the ones causing a change in the GC content. The blue arrows were manually added to show the positions of the two primer sets designed on *wzi* in this work.

### High-Resolution Melting analysis

We performed HRM experiments using ten primer sets on two strain collections. Four out of the ten primer sets were newly designed in this work (see above), while the remaining six were already available in literature^7^. The two strain collections were: i) the “background” collection, which includes 17 *K. pneumoniae* strains belonging to 17 different Sequence Types (STs); ii) the “outbreak” collection, which includes 11 *K. pneumoniae* strains isolated during a nosocomial outbreak. The obtained melting temperatures (“Tm”) of the three HRM replicates and their relative average temperature (“aTm”) values are reported in <u>Supplementary Table S1</u>.

### Primer sets and schemes comparison

For each of the ten primer sets we calculated the strain distance matrix among the background strains based on the aTm values (see Methods). The calculated aTm distances ranged from zero to three degrees, and the median distances varied among the genes (as shown in Figure 2). In particular, the two *wzi* primer sets showed median distance values significantly higher than those obtained for many of the other primer sets (see <u>Supplementary Table S2</u> for details).

**Figure 2.**
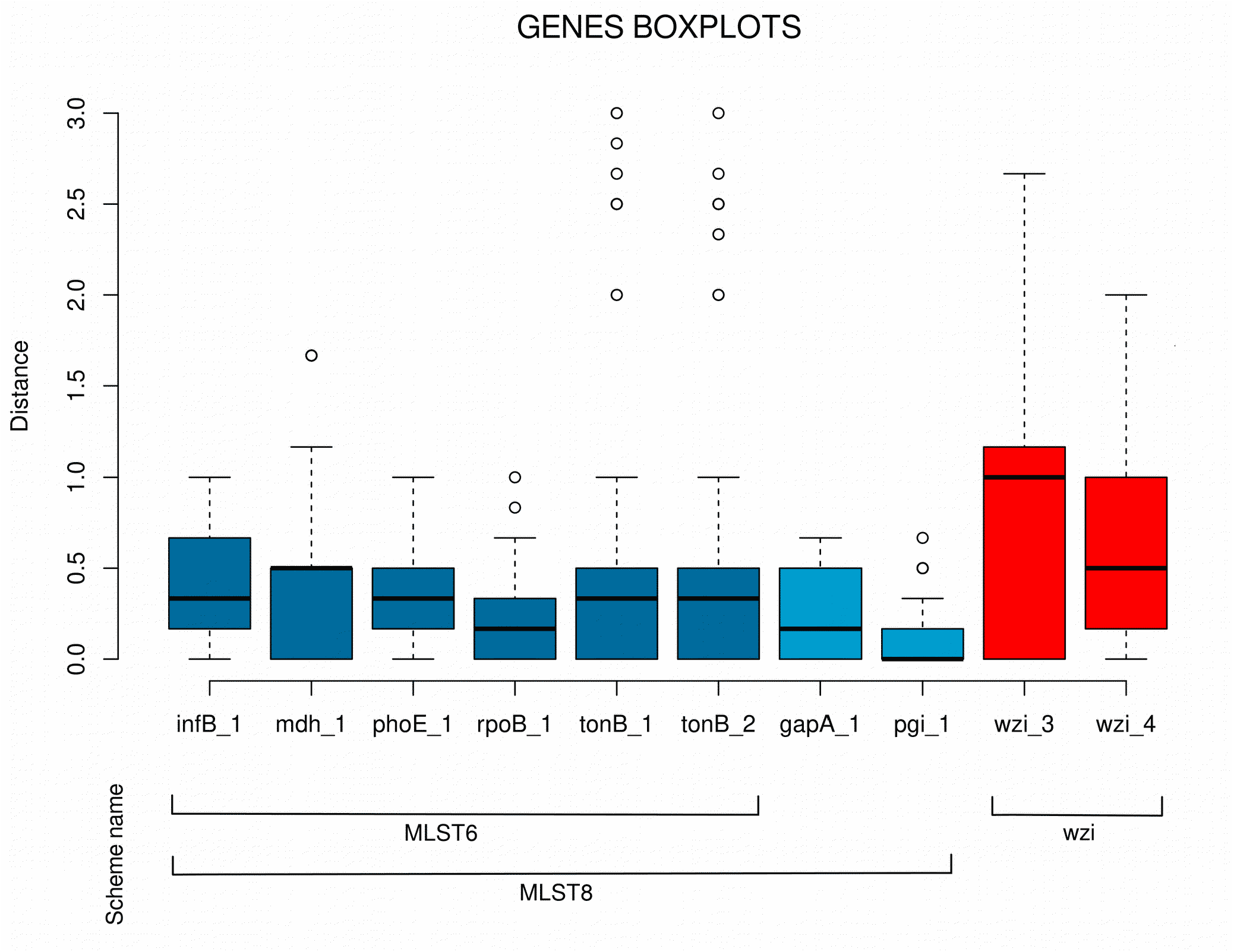
Distribution of the average melting temperature differences among the 17 *Klebsiella pneumoniae* strains for the ten primer sets. Boxes are the 25th and 75th quartiles divided by the medians, whiskers are 1.5x the interquartile ranges and dots are outliers. The lines in the bottom show the composition of the three primer schemes used in this work.

We also compared the aTm distance matrices of the following schemes:

- “MLST6”: which includes the HRM-MLST primer sets already present in literature;
- “MLST8”: which includes the MLST6 primer sets and the two newly designed;
- “wzi”: which includes the two primer sets designed for the *wzi* gene.

The median pairwise distance did not significantly change among the three schemes (Wilcoxon test with Holm post-hoc correction, p-value > 0.05) and the relative boxplot graphs are reported in <u>Supplementary Fig. S3</u>.

Furthermore, we compared the aTm distance matrices of wzi and MLST8 schemes for each strain pair of the background collection, subtracting the two matrices (see Figure 3). We found that, among all the 136 possible strain pairs, 66 (48.5%) showed a higher distance for wzi than MLST8. More in detail, the ST147 resulted better discriminated by MLST8 scheme, while the ST15 and ST307 resulted discriminated by wzi.

**Figure 3.**
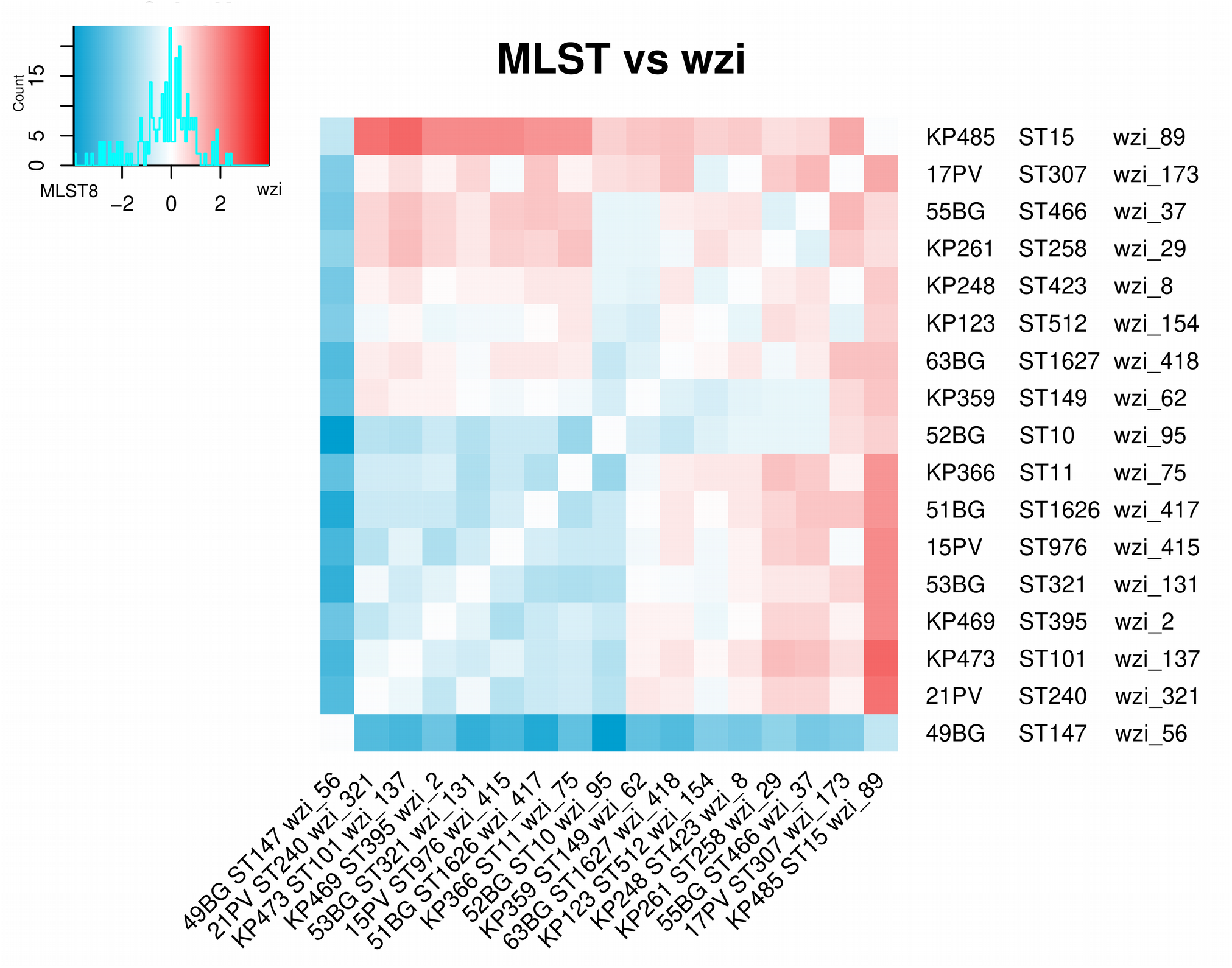
Arithmetic difference between the average melting temperature distance matrices computed among the 17 *Klebsiella pneumoniae* strains (selected to belong to 17 different STs) using the MLST8 scheme (eight primer sets on seven genes) and wzi scheme (two primer set on one gene). The heatmap colours range from blue to white to red: if the temperature distance between two strains is greater for the MLST8 than the wzi scheme the relative position on the heatmap is coloured in blue, otherwise in red.

### Whole Genome Sequencing-based strain typing

Genomic reads of the 11 *K. pneumoniae* strains of the outbreak collection and genomic reads of the eight strains of the background collection isolated during the San Raffaele hospital surveillance program were obtained.

Ten out of the 11 outbreak isolates belonged to the ST512 while the isolate “*BG-Kpn-22-18*” resulted to belong to the ST307. All the *wzi* alleles and the STs identified for the 28 *K. pneumoniae* strains (19 sequenced in this work and 9 from Gaiarsa and colleagues^10^) as well as their accession numbers are reported are reported in <u>Supplementary Table S3</u>.

### WGS-based outbreak reconstruction

An alignment of 66 core-SNPs was obtained from the 11 outbreak strains. The relative Maximum Likelihood phylogenetic tree is reported in <u>Supplementary Fig. S4</u>. The ST512 strains have SNP distances ranging from zero to four SNPs. Conversely, the SNP distances among the ST512 strains and the ST307 strain ranged from 63 to 66 SNPs.

### HRM-based outbreak reconstruction

The three dendrograms obtained by hierarchical clustering on the aTms strain distances for the schemes MLST6, MLST8 and wzi are reported in <u>Supplementary Fig. S5</u>, <u>Supplementary Fig. S6</u> and Figure 4, respectively. All the schemes correctly discriminated the outbreak ST512 strains from the ST307 one. Notably, only the wzi scheme correctly clustered the outbreak strains with the background strain of the same ST.

**Figure 4.**
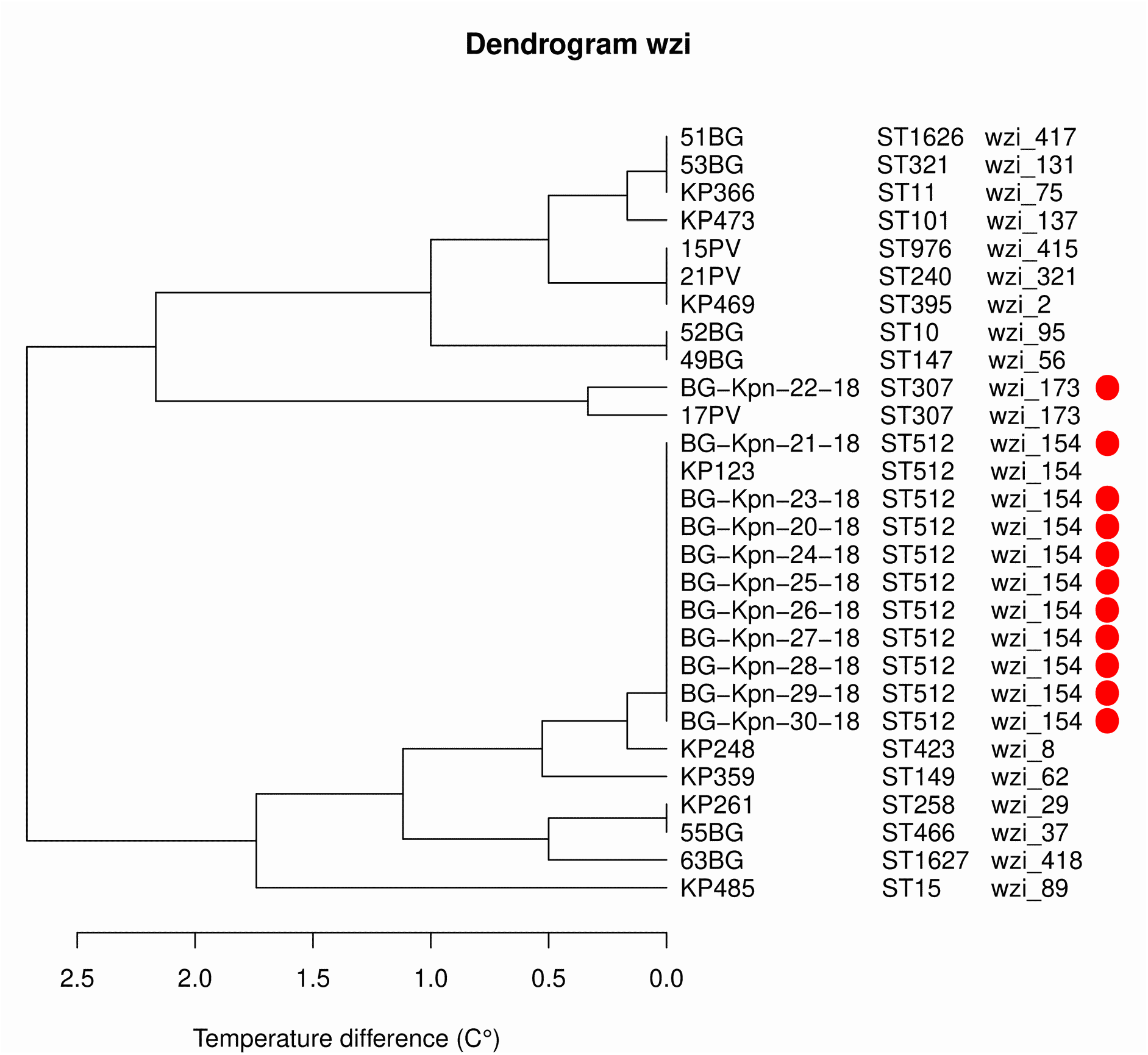
Dendrogram of the hierarchical clustering analysis on the average temperature distance matrix obtained using the wzi scheme. The 17 “background” strains belonging to 17 different MLSTs are written in black, while the 11 strains isolated during the nosocomial outbreak are highlighted by a red dot.

## Discussion

High Resolution Melting (HRM) is a real-time PCR analysis for the detection of mutations and polymorphisms^3,4^, also applicable for fast bacterial typing in hospital surveillance and real-time nosocomial outbreak detection^5^. Several works applied HRM to bacterial typing, exploiting Multi Locus Sequence Type (MLST) genes^7,11,12^, which has been considered the gold standard genes for bacterial typing for almost 20 years^13^. These genes have been selected to be housekeeping therefore they display low variability. In this work we show that it is possible to increase HRM discriminatory power using hypervariable genes.

On the other hand, the identification of the regions suitable for primer design can be challenging when the number of aligned sequences is high or when the gene is hypervariable. Thus, we developed EasyPrimer, a tool for the identification of the best regions for primer design for both HRM analysis and, more in general, for any kind of pan-PCR study. EasyPrimer shows, with an easy-to-read graphical output, which are the best regions for primer design: two conserved regions flanking a highly variable one. The on-line and the standalone versions of the tool are freely available at https://skynet.unimi.it/index.php/tools/.

We validated the tool designing HRM primers for the nosocomial pathogen *Klebsiella pneumoniae*. A scheme including six HRM primer sets for five out of the seven *K. pneumoniae* MLST genes was already available in literature^7^ (MLST6 scheme). Thus, we used EasyPrimer to design the primers for the remaining two MLST genes (*pgi* and *gapA*), obtaining a larger scheme with eight primer sets (MLST8 scheme). Furthermore, we designed two HRM primer sets for the hypervariable capsular gene *wzi*. We tested the discriminatory power of these schemes on 17 *K. pneumoniae* strains belonging to 17 different STs (see Methods) and we used the HRM approach to study an outbreak occurred in an Italian hospital.

Our analyses showed a good discriminatory power for both the MLST-based and *wzi*-based HRM assays. Indeed, both schemes successfully discriminated the 17 *K. pneumoniae* STs. Additionally, we want to highlight that the observed HRM discriminatory power was obtained using a BioRad CFX Connect real-time PCR instrument (BioRad, Hercules, California): a machine not specifically designed for HRM experiments but for real-time PCR, with a melting temperature sensitivity of 0.5°C (higher than other available HRM machines).

We found that the discriminatory power of an HRM scheme does not strictly depend on the number of genes but also on the genetic variability of the genes. Indeed, comparing the MLST6 and MLST8 schemes, we found that the median distance among the strains did not change significantly. As shown in Figure 4, wzi scheme can discriminate the highly epidemiologically relevant ST258, ST512 and ST307. Furthermore, the ST258 is better discriminated from ST307 and ST512 by the wzi scheme rather than MLST8 scheme (Figure 3). The wzi scheme contains two primer sets and this reduces drastically the amount of time and costs required for typing. For instance, using only two primer sets on a 96-well PCR plate, it is possible to type 15 isolates per run (five hours, including DNA extraction, HRM run and analysis of results) with a cost of ~5 euros each. This makes the HRM a feasible method for real-time surveillance and for a preliminary typing step in large epidemiological studies.

We applied the wzi scheme to the reconstruction of a nosocomial outbreak occurred in an Italian hospital. During the outbreak, 11 patients resulted colonized or infected by *K. pneumoniae* and the WGS typing revealed that the isolates belonged to two different clones. These clones were identified on the basis of core SNP distance (SNP distance < 5) and MLST profile (one isolate belongs to the ST307 and ten isolates to the ST512). As shown in Figure 4, the wzi scheme not only correctly discriminated the outbreak isolates of the two clones but it clustered them with the background isolates of the corresponding ST profile.

During the last years, WGS has revolutionized clinical microbiology, allowing the precise description of bacterial genomic features in few days (including the presence of resistance and/or virulence factors). Despite this, its application during real-time outbreak reconstructions still shows some limits: the time required to be completed, the cost and the necessity of qualified personnel for library preparation, bioinformatic analyses and results interpretation. Indeed, the complete sequencing of a bacterial strain genome costs at least ~100 euros (using an Illumina MiSeq machine) and requires one or two days for library preparation and 5-36 hours for sequencing. During the first days of a nosocomial outbreak the number of cases still increases slowly. In this time frame, it is crucial to quickly understand if the bacterial strains are genetically related, and if the clone is spreading in the nosocomial environment. In this situation HRM is a “first-line” typing technology to figure out when an outbreak is starting. Indeed, HRM is less precise than WGS but it can be reliable for a fast, preliminary bacterial typing, fundamental in the first days of a nosocomial outbreak. If the outbreak is identified, WGS could be used to further investigate the transmission dynamics Indeed, HRM assay represents a fast, simple and time/cost saving approach for bacterial typing, allowing to analyse several bacterial samples per days. Furthermore, this technique does not require advanced skills in molecular biology and the results can be analysed without the use of any specific software. This method can be useful also in veterinary and dairy farming settings: *K. pneumoniae* is a relevant veterinary pathogen and one of the most frequent cause of mastitis in dairy cattle^9^.

The use of hypervariable genes in HRM-based bacterial typing, such as *wzi* in *K. pneumoniae*, can drastically increase the discriminatory power of the method. With the large number of genomes available in databases it is now possible to find the most variable genes for a species. Unfortunately, it is not easy to identify the best regions to design primers in such hypervariable genes, particularly when hundreds of different alleles are available. EasyPrimer can represent a useful tool to overcome this limit.

## Methods

### Isolates collection

We considered two strain collections: the “background” and the “outbreak” collections. The background collection includes 17 strains belonging to 17 different STs: nine retrieved from a previously WGS typed^10^ bacterial collection (see <u>Supplementary Table S3</u> for details) and eight isolated at the San Raffaele hospital, Milan, during a WGS-based surveillance project (for details see <u>Supplementary Table S3</u>). The outbreak collection includes 11 *K. pneumoniae* isolates gathered during a 16 days nosocomial outbreak occurred in April 2018, in the Papa Giovanni XXIII hospital (Bergamo) (For details see <u>Supplementary Table S4</u>).

### DNA extraction and Whole-Genome Sequencing

The genomic DNA of the 11 outbreak strains was extracted using the DNeasy blood and tissue kit (Qiagen, Hilden, Germany) following the manufacturer’s instructions. The extracted DNA was sequenced using the Illumina Miseq platform with a 2 x 250 bp paired-end run, after Nextera XT library preparation. The genomic DNA of the eight strains isolated during the San Raffaele hospital surveillance was extracted using Maxwell 16 Cell DNA purification kit. The extracted DNA was sequenced using the NextSeq500 platform with 2 × 150 bp paired-ends runs, after Nextera XT library preparation. The genomic DNA of the nine strains isolated by Gaiarsa and colleagues^10^ were extracted using QIAsymphony Virus/Pathogen minikit, version 1 (Qiagen, Hilden, Germany) with the automated instrument QIAsimphony (Qiagen, Hilden, Germany) according to manufacturer’s instructions.

### High Resolution Melting primer design using EasyPrimer

The EasyPrimer tool was developed for the identification of the most suitable regions for primers design in HRM and, more in general, in pan-PCR experiments. Briefly, the tool starts from the gene alignment, evaluates the amount of genetic variation for each position and identifies the most reliable regions for primer design. EasyPrimer flags as good candidates for primer design two conserved regions flanking a highly variable one (taking into consideration, in advance, the optimal lengths of primers and amplicon). The user can decide either to evaluate the variability of the amplicon considering HRM-detectable SNPs only (the best option for HRM primer design) or all the SNPs (the best set for pan-PCR experiments). A detailed description of the algorithm is reported in the <u>Supplementary Note S1</u>.

To develop an HRM-based protocol for *K. pneumoniae* typing, we focused on the seven MLST genes and on the hypervariable capsular gene *wzi*^14^. The HRM primer sets for five out of the seven *K. pneumoniae* MLST genes were already available in literature^7^ (*infB, mdh, phoE, rpoB* and two sets on *tonB*). For the remaining two MLST genes (*pgi* and *gapA*) and for the *wzi* capsular gene (two primer sets) the primers were designed using EasyPrimer. For each gene the sequences were downloaded from the BigsDB database (https://bigsdb.pasteur.fr, 218 alleles for *pgi*, 183 for *gapA* and 563 for *wzi*), EasyPrimer was run and primer sets were designed on the basis of its output.

### High-Resolution Melting assays

We performed HRM assays using the genomic DNA extracted from each of the 28 *K. pneumoniae* strains included in this work, using each of the ten primer sets mentioned above. HRM analyses were performed on the BioRad CFX Connect real-time PCR System (BioRad, Hercules, California). Each 10µl reaction contained: 5µl of iTaq™ Universal SYBR® Green Supermix (BioRad, Hercules, California), 0.4µl of each primer (0.4µM) and 1µl of template DNA (25-50ng/µl). The thermal profile was as follows: 98°C for 2min, 40 cycles of [95°C for 7s, 61°C for 7s, and 72°C for 15s], 95°C for 2min, followed by HRM ramping from 70–95°C with fluorescence data acquisition at 0.5°C increments. Three technical replicates were performed for each strain and for each gene analysed. Negative controls were added in every run and for each gene.

### Comparison of the HRM primer sets and schemes

We compared the discriminatory power of the ten HRM primer sets on the 17 strains of the background collection. For each set we calculated the average melting temperatures (aTms) of the three replicates for each strain and the relative strain distance matrix based on the obtained aTms. Thus, we compared the discriminatory power of the different primer sets by comparing the relative distance matrix values using Wilcoxon test with Holm post-hoc correction.

Furthermore, we grouped the primers sets in three schemes (MLST6, MLST8 and wzi) and we compared the relative strain distance matrices using Wilcoxon test with Holm post-hoc correction. The scheme compositions were as follows: the MLST6 scheme included the six primer sets proposed by Andersson and colleagues^7^ for five MLST genes (with two primer sets for *tonB*); the MLST8 included all the MLST6 primer sets, the primers for *pgi* and *gapA* (newly designed in this work using the EasyPrimer tool); the wzi scheme included the two primer sets for the *wzi* gene (newly designed in this work). For details see Table 1 and Figure 2.

Then, we compared the discriminatory power of MLST8 and wzi schemes by subtracting the relative distance matrices (wzi – MLST8) and studying the obtained matrix with heatmaps.

All these analyses were performed using R (https://www.r-project.org/) and the R libraries Ape and Gplots.

### HRM-based outbreak reconstruction

From the aTms of the outbreak and background collections we calculated the distance matrices for MLST6, MLST8 and wzi primer schemes (for more details see above) and clustered the strains using the hierarchical clustering method implemented in the Hclust function in R.

### WGS-based strain typing

We retrieved the reads of the nine *K. pneumoniae* strains previously WGS-typed by Gaiarsa and colleagues^10^ from NCBI database using fastq-dump tool (for accession numbers see <u>Supplementary Table S3</u>).

Then we performed *de novo* assembly on the reads obtained from the 19 strains sequenced in this work and those from the nine strains from database using SPAdes software^15^.

We retrieved the *wzi* allele sequences from BigsDB database and we annotated the *wzi* allele present in each of the 28 genome assemblies included in the study by Blastn search and manual curation of the results.

We retrieved the sequences of the *K. pneumoniae* MLST gene alleles and the relative scheme tables from the BigsDB database. Thus, we determined the MLST profiles using an in-house Blastn-based Perl script.

### Core-SNP-based phylogenetic reconstruction on outbreak strains

We aligned the reads obtained from the 11 outbreak strains against the NTUH_K2044 reference genome (accession number NC_016845.1), and performed the SNPs calling following the GATK best practice procedure. We masked SNPs localized within repeated regions, identified using MUMmer^16^, or prophages, identified using PhiSpy^17^, and we called the core-SNPs among the strains using an in-house Python script. Thus, we subjected the core-SNPs alignment to phylogenetic analysis as follows: the best evolutionary model was assessed by ModelTest-NG and phylogenetic reconstruction was performed using the selected best model, with RAxML8 software^18^. We evaluated the core-SNPs distances among the strains using the R Ape library.

### Data Availability Statement

We deposited all Illumina sequence data from the 19 strains in NCBI’s Short Read Archive under BioProject ID *(pending)* and all Illumina data were deposited under BioProject ID (*pending*)

## Acknowledgements

Thanks to the Romeo ed Enrica Invernizzi Foundation and to professor Claudio Bandi for supporting and revising the project.

## Author Contribution Statement

MP developed the tool, performed the HRM experiments and drafted the paper. AP performed the HRM experiments and revised the manuscript. SP designed the primers and revised the manuscript. DDC implemented the tool online. MC, FG, FV collected the samples and extracted the DNA. PM, DMC, CF collected the samples. GVZ wrote the paper. FC conceived and designed the experiments and wrote the paper. All authors read, revised and approved the final manuscript.

## Additional information

### Accession codes

*pending*

### Supplementary information

accompanies this paper at doi:

