## Supplemental material for "Fast *Klebsiella pneumoniae* typing for outbreak reconstruction: an highly discriminatory HRM protocol on *wzi* capsular gene developed using EasyPrimer tool"

### Supplementary Material

**Supplementary Note S1.** The EasyPrimer algorithm.

EasyPrimer is a Python-R pipeline to assist the primer design procedure. it was developed to identify the best regions to design primers in an alignment. Indeed, the tool finds the best “primer-amplicon-primer” gene region, where two highly conserved gene regions (suggested for primer design) flank a highly variable region (the “amplicon region”).

The algorithm scans the alignment to find - at least - one highly variable region (amplicon region) flanked by two conserved regions (primer regions). For the computation of the highly variable region, the user can decide to consider all the Single Nucleotide Polymorphism (SNPs) or only the HRM-detectable ones (HRM-SNP): a HRM-SNP is a SNP that confers a change in CG content (i.e., any A↔G or T↔G or A↔C or T↔G counts as a HRM-SNP). EasyPrimer will identify the most HRM discriminatory region in the alignment when HRM SNPs are used to score the variability of the amplicon region. Conversely, considering all the SNPs the tool will identify the most variable region in general. Additionally, the user provides length ranges both for primers and amplicon, the tool suggests regions for primer design with a reasonable spacing, meaning that they always amplify a region of optimal length.

The main EasyPrimer output is a graph (plotted in a PDF file) in which the alignment consensus sequence is reported on the x-axes and the information about the positions of the best primer/amplicon regions are represented with blue and red curves.

**The algorithm:**

The inputs of the tool are:

i) A Fasta sequence of the gene to be analysed;
ii) A minimum and a maximum nucleotide length allowed for the primers and the amplicon;
iii) A percentile threshold of variability for primers selection (see below);
iv) A percentile threshold of variability for amplicon selection (see below);
v) A consensus threshold value (see below);

The steps are the following:

1. Aligning the gene sequences using the MUSCLE software (MUltiple Sequence Comparison by Log- Expectation)[1]
2. Computing, for each position of the alignment, the frequencies of A, T, C, G and gaps
3. Computing for each position the information of the entropy using Shannon index on the basis of A, T, C, G and gaps frequencies as follows:

$$H' = [f_A \times \log(f_A)] + [f_T \times \log(f_T)] + [f_C \times \log(f_C)] + [f_G \times \log(f_G)] + [f_{gaps} \times \log(f_{gaps})]$$

4. Estimating the frequency of mutation among strains. If the user decided to use all the possible SNPs to evaluate the amplicon variability the score is calculated as follows:

$$f_{SNP} = \frac{(f_A \times f_G) + (f_T \times f_G) + (f_A \times f_C) + (f_T \times f_C) + (f_A \times f_T) + (f_G \times f_C)}{amplicon\ length}$$

Otherwise, if the user decided to work with HRM-SNPs only the formula becomes:

$$f_{HRM} = \frac{(f_A \times f_G) + (f_T \times f_G) + (f_A \times f_C) + (f_T \times f_C)}{amplicon\ length}$$

5. For each possible combination of primers and amplicon lengths a window composed by primer-amplicon-primer sections is generated and shifted on the sequence alignment. For each window position, the average H' is calculated for the primer regions and the average $f_{HRM}$ or $f_{SNP}$ is calculated for the amplicon region
6. Selection of the windows with total $H'$ below the primer percentile threshold in the primer region and $f_{HRM}/f_{SNP}$above the amplicon percentile threshold in the amplicon region.
7. The consensus of the alignment is obtained
8. The results are plotted on a graph in which the consensus sequence is reported on the x-axes and blue/red lines are obtained as follows: for each alignment position, the frequency of selected windows for which the position is included in the primer section is represented in blue, while the frequency for which the position in included in the amplicon section is represented in red

Notes on the consensus notation:
The consensus sequence is written the IUPAC notation for DNA both for the bases and the degenerated residues. some symbols and notation were added to give more information:

- Red gaps symbols (-) are added to indicate that gaps in the alignment are greater than consensus threshold (more than 5% in default) and less than 1 - consensus threshold (less than 95% in default).
- Asterics (*) indicate a gap present more than 1 - consensus threshold (more than 95% in defaults), this indicate an insertion in one or few variances in the gene. The user will probably want to consider that position as 'not existing' during the primer design process

The algorithm is summarized in the flow diagram in Figure S7.

**Table S1.** List of all the melting temperatures obtained during the HRM analyses for all the strains and for all the primer sets.

| Strain name | MLST profile | Gene name | Primer set name | Melting T 1 (°C) | Melting T 2 (°C) | Melting T 3 (°C) | Average melting T (°C) |
|---|---|---|---|---|---|---|---|
| 15PV | ST976 | *infB* | infB_1 | 84 | 84 | 84 | 84 |
| 15PV | ST976 | *mdh* | mdh_1 | 80.5 | 80.5 | 80.5 | 80.5 |
| 15PV | ST976 | *phoE* | phoE_1 | 80 | 80 | 80 | 80 |
| 15PV | ST976 | *rpoB* | rpoB_1 | 83.5 | 83.5 | 83 | 83.3 |
| 15PV | ST976 | *tonB* | tonB_1 | 86.5 | 86.5 | 86 | 86.3 |
| 15PV | ST976 | *tonB* | tonB_2 | 89 | 89 | 89 | 89 |
| 15PV | ST976 | *gapA* | gapA_1 | 84.5 | 84.5 | 84.5 | 84.5 |
| 15PV | ST976 | *pgi* | pgi_1 | 86 | 86 | 86 | 86 |
| 15PV | ST976 | *wzi* | wzi_3 | 85 | 85 | 85 | 85 |
| 15PV | ST976 | *wzi* | wzi_4 | 83.5 | 83.5 | 83.5 | 83.5 |
| 17PV | ST307 | *pgi* | pgi_1 | 86 | 85.5 | 85.5 | 85.7 |
| 17PV | ST307 | *wzi* | wzi_3 | 84.5 | 84 | 84 | 84.2 |
| 17PV | ST307 | *wzi* | wzi_4 | 82.5 | 82.5 | 82.5 | 82.5 |
| 17PV | ST307 | *infB* | infB_1 | 84 | 83.5 | 83.5 | 83.7 |
| 17PV | ST307 | *mdh* | mdh_1 | 80 | 80 | 80 | 80 |
| 17PV | ST307 | *phoE* | phoE_1 | 81 | 81 | 80.5 | 80.8 |
| 17PV | ST307 | *rpoB* | rpoB_1 | 83.5 | 83.5 | 83.5 | 83.5 |
| 17PV | ST307 | *tonB* | tonB_1 | 86.5 | 86 | 86 | 86.2 |
| 17PV | ST307 | *tonB* | tonB_2 | 89.5 | 89.5 | 89.5 | 89.5 |
| 17PV | ST307 | *gapA* | gapA_1 | 85.5 | 85 | 85 | 85.2 |
| 21PV | ST307 | *infB* | infB_1 | 84.5 | 84.5 | 84 | 84.3 |
| 21PV | ST307 | *mdh* | mdh_1 | 80.5 | 80.5 | 80.5 | 80.5 |
| 21PV | ST307 | *phoE* | phoE_1 | 81 | 81 | 81 | 81 |
| 21PV | ST307 | *rpoB* | rpoB_1 | 83.5 | 83.5 | 83.5 | 83.5 |
| 21PV | ST307 | *tonB* | tonB_1 | 86.5 | 86.5 | 86.5 | 86.5 |
| 21PV | ST307 | *tonB* | tonB_2 | 89 | 89 | 89 | 89 |
| 21PV | ST307 | *gapA* | gapA_1 | 85 | 85 | 85 | 85 |
| 21PV | ST307 | *pgi* | pgi_1 | 86 | 86 | 86 | 86 |
| 21PV | ST307 | *wzi* | wzi_3 | 85 | 85 | 85 | 85 |
| 21PV | ST307 | *wzi* | wzi_4 | 83.5 | 83.5 | 83.5 | 83.5 |
| 49BG | ST147 | *infB* | infB_1 | 83.5 | 83.5 | 83.5 | 83.5 |
| 49BG | ST147 | *mdh* | mdh_1 | 79.5 | 80 | 79.5 | 79.7 |
| 49BG | ST147 | *phoE* | phoE_1 | 80.5 | 80.5 | 80.5 | 80.5 |
| 49BG | ST147 | *rpoB* | rpoB_1 | 83.5 | 83.5 | 84 | 83.7 |
| 49BG | ST147 | *tonB* | tonB_1 | 89 | 89 | 89 | 89 |
| 49BG | ST147 | *tonB* | tonB_2 | 86.5 | 86.5 | 86.5 | 86.5 |
| 49BG | ST147 | *gapA* | gapA_1 | 85.5 | 85 | 85 | 85.2 |
| 49BG | ST147 | *pgi* | pgi_1 | 86 | 86 | 86 | 86 |
| 49BG | ST147 | *wzi* | wzi_3 | 85 | 85 | 85 | 85 |
| 49BG | ST147 | *wzi* | wzi_4 | 84.5 | 84.5 | 84.5 | 84.5 |
| 51BG | ST1626 | *infB* | infB_1 | 83.5 | 83.5 | 83 | 83.3 |
| 51BG | ST1626 | *mdh* | mdh_1 | 80 | 80 | 80 | 80 |
| 51BG | ST1626 | *phoE* | phoE_1 | 80.5 | 80.5 | 80.5 | 80.5 |

| Strain name | MLST profile | Gene name | Primer set name | Melting T 1 (°C) | Melting T 2 (°C) | Melting T 3 (°C) | Average melting T (°C) |
|---|---|---|---|---|---|---|---|
| 51BG | ST1626 | *rpoB* | rpoB_1 | 83.5 | 83.5 | 83.5 | 83.5 |
| 51BG | ST1626 | *tonB* | tonB_1 | 86 | 86 | 86 | 86 |
| 51BG | ST1626 | *tonB* | tonB_2 | 89 | 89 | 89 | 89 |
| 51BG | ST1626 | *gapA* | gapA_1 | 85 | 85 | 85 | 85 |
| 51BG | ST1626 | *pgi* | pgi_1 | 86 | 86 | 86 | 86 |
| 51BG | ST1626 | *wzi* | wzi_3 | 85 | 85 | 85 | 85 |
| 51BG | ST1626 | *wzi* | wzi_4 | 84 | 84 | 84 | 84 |
| 52BG | ST10 | *infB* | infB_1 | 83.5 | 83.5 | 83.5 | 83.5 |
| 52BG | ST10 | *mdh* | mdh_1 | 78.5 | 79 | 79 | 78.8 |
| 52BG | ST10 | *phoE* | phoE_1 | 80 | 80.5 | 80 | 80.2 |
| 52BG | ST10 | *rpoB* | rpoB_1 | 83 | 83 | 83.5 | 83.2 |
| 52BG | ST10 | *tonB* | tonB_1 | 86 | 86 | 86 | 86 |
| 52BG | ST10 | *tonB* | tonB_2 | 88.5 | 89 | 89 | 88.8 |
| 52BG | ST10 | *gapA* | gapA_1 | 84.5 | 85 | 84.5 | 84.7 |
| 52BG | ST10 | *pgi* | pgi_1 | 86 | 86 | 86 | 86 |
| 52BG | ST10 | *wzi* | wzi_3 | 85 | 85 | 85 | 85 |
| 52BG | ST10 | *wzi* | wzi_4 | 84.5 | 84.5 | 84.5 | 84.5 |
| 53BG | ST321 | *infB* | infB_1 | 84 | 84.5 | 84 | 84.2 |
| 53BG | ST321 | *mdh* | mdh_1 | 80 | 80 | 80 | 80 |
| 53BG | ST321 | *phoE* | phoE_1 | 81 | 81 | 81 | 81 |
| 53BG | ST321 | *rpoB* | rpoB_1 | 83 | 83 | 83 | 83 |
| 53BG | ST321 | *tonB* | tonB_1 | 86.5 | 86.5 | 86.5 | 86.5 |
| 53BG | ST321 | *tonB* | tonB_2 | 89 | 89 | 89 | 89 |
| 53BG | ST321 | *gapA* | gapA_1 | 85 | 85 | 85 | 85 |
| 53BG | ST321 | *pgi* | pgi_1 | 86 | 86 | 86 | 86 |
| 53BG | ST321 | *wzi* | wzi_3 | 85 | 85 | 85 | 85 |
| 53BG | ST321 | *wzi* | wzi_4 | 84 | 84 | 84 | 84 |
| 55BG | ST466 | *infB* | infB_1 | 83.5 | 83.5 | 83 | 83.3 |
| 55BG | ST466 | *mdh* | mdh_1 | 80.5 | 80.5 | 80.5 | 80.5 |
| 55BG | ST466 | *phoE* | phoE_1 | 80.5 | 80.5 | 80.5 | 80.5 |
| 55BG | ST466 | *rpoB* | rpoB_1 | 83.5 | 83.5 | 83.5 | 83.5 |
| 55BG | ST466 | *tonB* | tonB_1 | 86.5 | 86.5 | 86.5 | 86.5 |
| 55BG | ST466 | *tonB* | tonB_2 | 89 | 89 | 89 | 89 |
| 55BG | ST466 | *gapA* | gapA_1 | 85 | 85 | 85 | 85 |
| 55BG | ST466 | *pgi* | pgi_1 | 86 | 86 | 86 | 86 |
| 55BG | ST466 | *wzi* | wzi_3 | 83.5 | 83.5 | 83.5 | 83.5 |
| 55BG | ST466 | *wzi* | wzi_4 | 84.5 | 84.5 | 84.5 | 84.5 |
| 63BG | ST1627 | *infB* | infB_1 | 83.5 | 83.5 | 83 | 83.3 |
| 63BG | ST1627 | *mdh* | mdh_1 | 80.5 | 80.5 | 80.5 | 80.5 |
| 63BG | ST1627 | *phoE* | phoE_1 | 80.5 | 80.5 | 80.5 | 80.5 |
| 63BG | ST1627 | *rpoB* | rpoB_1 | 83.5 | 83.5 | 83.5 | 83.5 |
| 63BG | ST1627 | *tonB* | tonB_1 | 86.5 | 86.5 | 86.5 | 86.5 |
| 63BG | ST1627 | *tonB* | tonB_2 | 88.5 | 89.5 | 89.5 | 89.2 |
| 63BG | ST1627 | *gapA* | gapA_1 | 85 | 85 | 85 | 85 |
| 63BG | ST1627 | *pgi* | pgi_1 | 86 | 86 | 86.5 | 86.2 |
| 63BG | ST1627 | *wzi* | wzi_3 | 84 | 84 | 84 | 84 |

| Strain name | MLST profile | Gene name | Primer set name | Melting T 1 (°C) | Melting T 2 (°C) | Melting T 3 (°C) | Average melting T (°C) |
|---|---|---|---|---|---|---|---|
| 63BG | ST1627 | *wzi* | wzi_4 | 84.5 | 84.5 | 84.5 | 84.5 |
| BG-Kpn-20-18 | ST512 | *wzi* | wzi_3 | 84 | 84 | 84 | 84 |
| BG-Kpn-20-18 | ST512 | *wzi* | wzi_4 | 83.5 | 83.5 | 83.5 | 83.5 |
| BG-Kpn-20-18 | ST512 | *pgi* | pgi_1 | 86 | 86 | 86 | 86 |
| BG-Kpn-20-18 | ST512 | *infB* | infB_1 | 83.5 | 83.5 | 83.5 | 83.5 |
| BG-Kpn-20-18 | ST512 | *mdh* | mdh_1 | 80.5 | 80.5 | 80.5 | 80.5 |
| BG-Kpn-20-18 | ST512 | *phoE* | phoE_1 | 80.5 | 80.5 | 80.5 | 80.5 |
| BG-Kpn-20-18 | ST512 | *rpoB* | rpoB_1 | 83.5 | 83.5 | 83 | 83.3 |
| BG-Kpn-20-18 | ST512 | *tonB* | tonB_1 | 86.5 | 86.5 | 86.5 | 86.5 |
| BG-Kpn-20-18 | ST512 | *tonB* | tonB_2 | 88.5 | 88.5 | 88.5 | 88.5 |
| BG-Kpn-20-18 | ST512 | *gapA* | gapA_1 | 85 | 84.5 | 84.5 | 84.7 |
| BG-Kpn-21-18 | ST512 | *pgi* | pgi_1 | 86.5 | 86 | 86 | 86.2 |
| BG-Kpn-21-18 | ST512 | *wzi* | wzi_3 | 84 | 84 | 84 | 84 |
| BG-Kpn-21-18 | ST512 | *wzi* | wzi_4 | 83.5 | 83.5 | 83.5 | 83.5 |
| BG-Kpn-21-18 | ST512 | *infB* | infB_1 | 83.5 | 83.5 | 83.5 | 83.5 |
| BG-Kpn-21-18 | ST512 | *mdh* | mdh_1 | 80.5 | 80.5 | 80.5 | 80.5 |
| BG-Kpn-21-18 | ST512 | *phoE* | phoE_1 | 80.5 | 80.5 | 80.5 | 80.5 |
| BG-Kpn-21-18 | ST512 | *rpoB* | rpoB_1 | 83.5 | 83.5 | 83.5 | 83.5 |
| BG-Kpn-21-18 | ST512 | *tonB* | tonB_1 | 86.5 | 86.5 | 86.5 | 86.5 |
| BG-Kpn-21-18 | ST512 | *tonB* | tonB_2 | 88.5 | 88.5 | 88.5 | 88.5 |
| BG-Kpn-21-18 | ST512 | *gapA* | gapA_1 | 85 | 85 | 85 | 85 |
| BG-Kpn-22-18 | ST307 | *pgi* | pgi_1 | 86 | 86 | 86 | 86 |
| BG-Kpn-22-18 | ST307 | *wzi* | wzi_3 | 84.5 | 84.5 | 84.5 | 84.5 |
| BG-Kpn-22-18 | ST307 | *wzi* | wzi_4 | 82.5 | 82.5 | 82.5 | 82.5 |
| BG-Kpn-22-18 | ST307 | *infB* | infB_1 | 84 | 84 | 84 | 84 |
| BG-Kpn-22-18 | ST307 | *mdh* | mdh_1 | 79.5 | 79.5 | 79.5 | 79.5 |
| BG-Kpn-22-18 | ST307 | *phoE* | phoE_1 | 80.5 | 80.5 | 80.5 | 80.5 |
| BG-Kpn-22-18 | ST307 | *rpoB* | rpoB_1 | 83.5 | 83.5 | 83 | 83.3 |
| BG-Kpn-22-18 | ST307 | *tonB* | tonB_1 | 86.5 | 86.5 | 86.5 | 86.5 |
| BG-Kpn-22-18 | ST307 | *tonB* | tonB_2 | 89 | 89 | 89 | 89 |
| BG-Kpn-22-18 | ST307 | *gapA* | gapA_1 | 85 | 85 | 85 | 85 |
| BG-Kpn-23-18 | ST512 | *pgi* | pgi_1 | 86 | 86 | 86.5 | 86.2 |
| BG-Kpn-23-18 | ST512 | *wzi* | wzi_3 | 84 | 84 | 84 | 84 |
| BG-Kpn-23-18 | ST512 | *wzi* | wzi_4 | 83.5 | 83.5 | 83.5 | 83.5 |
| BG-Kpn-23-18 | ST512 | *infB* | infB_1 | 83 | 83 | 83 | 83 |
| BG-Kpn-23-18 | ST512 | *mdh* | mdh_1 | 80.5 | 80.5 | 80.5 | 80.5 |
| BG-Kpn-23-18 | ST512 | *phoE* | phoE_1 | 80.5 | 80.5 | 80.5 | 80.5 |
| BG-Kpn-23-18 | ST512 | *rpoB* | rpoB_1 | 83.5 | 83.5 | 83.5 | 83.5 |
| BG-Kpn-23-18 | ST512 | *tonB* | tonB_1 | 86.5 | 86.5 | 86.5 | 86.5 |
| BG-Kpn-23-18 | ST512 | *tonB* | tonB_2 | 88.5 | 88.5 | 88.5 | 88.5 |
| BG-Kpn-23-18 | ST512 | *gapA* | gapA_1 | 84.5 | 84.5 | 84.5 | 84.5 |
| BG-Kpn-24-18 | ST512 | *pgi* | pgi_1 | 86.5 | 86 | 86 | 86.2 |
| BG-Kpn-24-18 | ST512 | *wzi* | wzi_3 | 84 | 84 | 84 | 84 |
| BG-Kpn-24-18 | ST512 | *wzi* | wzi_4 | 83.5 | 83.5 | 83.5 | 83.5 |
| BG-Kpn-24-18 | ST512 | *infB* | infB_1 | 83.5 | 83.5 | 83.5 | 83.5 |
| BG-Kpn-24-18 | ST512 | *mdh* | mdh_1 | 80.5 | 80.5 | 80.5 | 80.5 |

| Strain name | MLST profile | Gene name | Primer set name | Melting T 1 (°C) | Melting T 2 (°C) | Melting T 3 (°C) | Average melting T (°C) |
|---|---|---|---|---|---|---|---|
| BG-Kpn-24-18 | ST512 | *phoE* | phoE_1 | 81 | 81 | 81 | 81 |
| BG-Kpn-24-18 | ST512 | *rpoB* | rpoB_1 | 83.5 | 83.5 | 83.5 | 83.5 |
| BG-Kpn-24-18 | ST512 | *tonB* | tonB_1 | 86.5 | 86.5 | 86.5 | 86.5 |
| BG-Kpn-24-18 | ST512 | *tonB* | tonB_2 | 89 | 89 | 89 | 89 |
| BG-Kpn-24-18 | ST512 | *gapA* | gapA_1 | 84.5 | 84.5 | 84.5 | 84.5 |
| BG-Kpn-25-18 | ST512 | *pgi* | pgi_1 | 86 | 86 | 86 | 86 |
| BG-Kpn-25-18 | ST512 | *wzi* | wzi_3 | 84 | 84 | 84 | 84 |
| BG-Kpn-25-18 | ST512 | *wzi* | wzi_4 | 83.5 | 83.5 | 83.5 | 83.5 |
| BG-Kpn-25-18 | ST512 | *infB* | infB_1 | 83.5 | 83.5 | 83.5 | 83.5 |
| BG-Kpn-25-18 | ST512 | *mdh* | mdh_1 | 80.5 | 80.5 | 80.5 | 80.5 |
| BG-Kpn-25-18 | ST512 | *phoE* | phoE_1 | 81 | 81 | 81 | 81 |
| BG-Kpn-25-18 | ST512 | *rpoB* | rpoB_1 | 83.5 | 83.5 | 83.5 | 83.5 |
| BG-Kpn-25-18 | ST512 | *tonB* | tonB_1 | 86.5 | 86.5 | 86.5 | 86.5 |
| BG-Kpn-25-18 | ST512 | *tonB* | tonB_2 | 88.5 | 88.5 | 88.5 | 88.5 |
| BG-Kpn-25-18 | ST512 | *gapA* | gapA_1 | 84.5 | 84.5 | 84.5 | 84.5 |
| BG-Kpn-26-18 | ST512 | *pgi* | pgi_1 | 86 | 86 | 86 | 86 |
| BG-Kpn-26-18 | ST512 | *wzi* | wzi_3 | 84 | 84 | 84 | 84 |
| BG-Kpn-26-18 | ST512 | *wzi* | wzi_4 | 83.5 | 83.5 | 83.5 | 83.5 |
| BG-Kpn-26-18 | ST512 | *infB* | infB_1 | 83.5 | 83.5 | 83.5 | 83.5 |
| BG-Kpn-26-18 | ST512 | *mdh* | mdh_1 | 80.5 | 80.5 | 80.5 | 80.5 |
| BG-Kpn-26-18 | ST512 | *phoE* | phoE_1 | 81 | 81 | 80.5 | 80.8 |
| BG-Kpn-26-18 | ST512 | *rpoB* | rpoB_1 | 83.5 | 83.5 | 83.5 | 83.5 |
| BG-Kpn-26-18 | ST512 | *tonB* | tonB_1 | 86.5 | 86.5 | 86.5 | 86.5 |
| BG-Kpn-26-18 | ST512 | *tonB* | tonB_2 | 88.5 | 88.5 | 88.5 | 88.5 |
| BG-Kpn-26-18 | ST512 | *gapA* | gapA_1 | 84.5 | 84.5 | 84.5 | 84.5 |
| BG-Kpn-27-18 | ST512 | *pgi* | pgi_1 | 86 | 86 | 86 | 86 |
| BG-Kpn-27-18 | ST512 | *wzi* | wzi_3 | 84 | 84 | 84 | 84 |
| BG-Kpn-27-18 | ST512 | *wzi* | wzi_4 | 83.5 | 83.5 | 83.5 | 83.5 |
| BG-Kpn-27-18 | ST512 | *infB* | infB_1 | 83.5 | 83.5 | 83.5 | 83.5 |
| BG-Kpn-27-18 | ST512 | *mdh* | mdh_1 | 80.5 | 80.5 | 80.5 | 80.5 |
| BG-Kpn-27-18 | ST512 | *phoE* | phoE_1 | 81 | 81 | 81 | 81 |
| BG-Kpn-27-18 | ST512 | *rpoB* | rpoB_1 | 83.5 | 83.5 | 83.5 | 83.5 |
| BG-Kpn-27-18 | ST512 | *tonB* | tonB_1 | 86.5 | 86.5 | 86.5 | 86.5 |
| BG-Kpn-27-18 | ST512 | *tonB* | tonB_2 | 88.5 | 88.5 | 89 | 88.7 |
| BG-Kpn-27-18 | ST512 | *gapA* | gapA_1 | 84.5 | 84.5 | 84.5 | 84.5 |
| BG-Kpn-28-18 | ST512 | *pgi* | pgi_1 | 86 | 86 | 86 | 86 |
| BG-Kpn-28-18 | ST512 | *wzi* | wzi_3 | 84 | 84 | 84 | 84 |
| BG-Kpn-28-18 | ST512 | *wzi* | wzi_4 | 83.5 | 83.5 | 83.5 | 83.5 |
| BG-Kpn-28-18 | ST512 | *infB* | infB_1 | 83.5 | 83 | 83.5 | 83.3 |
| BG-Kpn-28-18 | ST512 | *mdh* | mdh_1 | 80.5 | 80.5 | 80.5 | 80.5 |
| BG-Kpn-28-18 | ST512 | *phoE* | phoE_1 | 81 | 81 | 81 | 81 |
| BG-Kpn-28-18 | ST512 | *rpoB* | rpoB_1 | 83.5 | 83.5 | 83.5 | 83.5 |
| BG-Kpn-28-18 | ST512 | *tonB* | tonB_1 | 86.5 | 86.5 | 86.5 | 86.5 |
| BG-Kpn-28-18 | ST512 | *tonB* | tonB_2 | 89 | 89 | 89 | 89 |
| BG-Kpn-28-18 | ST512 | *gapA* | gapA_1 | 84.5 | 84.5 | 84.5 | 84.5 |
| BG-Kpn-29-18 | ST512 | *pgi* | pgi_1 | 86 | 86 | 86 | 86 |

| Strain name | MLST profile | Gene name | Primer set name | Melting T 1 (°C) | Melting T 2 (°C) | Melting T 3 (°C) | Average melting T (°C) |
|---|---|---|---|---|---|---|---|
| BG-Kpn-29-18 | ST512 | *wzi* | wzi_3 | 84 | 84 | 84 | 84 |
| BG-Kpn-29-18 | ST512 | *wzi* | wzi_4 | 83.5 | 83.5 | 83.5 | 83.5 |
| BG-Kpn-29-18 | ST512 | *infB* | infB_1 | 83.5 | 83.5 | 83.5 | 83.5 |
| BG-Kpn-29-18 | ST512 | *mdh* | mdh_1 | 80.5 | 80.5 | 80.5 | 80.5 |
| BG-Kpn-29-18 | ST512 | *phoE* | phoE_1 | 81 | 81 | 81 | 81 |
| BG-Kpn-29-18 | ST512 | *rpoB* | rpoB_1 | 83.5 | 83.5 | 83.5 | 83.5 |
| BG-Kpn-29-18 | ST512 | *tonB* | tonB_1 | 86.5 | 86.5 | 86.5 | 86.5 |
| BG-Kpn-29-18 | ST512 | *gapA* | gapA_1 | 84.5 | 84.5 | 84.5 | 84.5 |
| BG-Kpn-29-18 | ST512 | *tonB* | tonB_2 | 89 | 89 | 89 | 89 |
| BG-Kpn-30-18 | ST512 | *pgi* | pgi_1 | 86 | 86 | 86 | 86 |
| BG-Kpn-30-18 | ST512 | *wzi* | wzi_3 | 84 | 84 | 84 | 84 |
| BG-Kpn-30-18 | ST512 | *wzi* | wzi_4 | 83.5 | 83.5 | 83.5 | 83.5 |
| BG-Kpn-30-18 | ST512 | *infB* | infB_1 | 83.5 | 83.5 | 83.5 | 83.5 |
| BG-Kpn-30-18 | ST512 | *mdh* | mdh_1 | 80.5 | 80.5 | 80.5 | 80.5 |
| BG-Kpn-30-18 | ST512 | *phoE* | phoE_1 | 81 | 81 | 81 | 81 |
| BG-Kpn-30-18 | ST512 | *rpoB* | rpoB_1 | 83.5 | 83.5 | 83.5 | 83.5 |
| BG-Kpn-30-18 | ST512 | *tonB* | tonB_1 | 86.5 | 86.5 | 86.5 | 86.5 |
| BG-Kpn-30-18 | ST512 | *tonB* | tonB_2 | 89 | 89 | 89 | 89 |
| BG-Kpn-30-18 | ST512 | *gapA* | gapA_1 | 84.5 | 84.5 | 84.5 | 84.5 |
| KP123 | ST512 | *infB* | infB_1 | 83.5 | 83.5 | 83 | 83.3 |
| KP123 | ST512 | *mdh* | mdh_1 | 80.5 | 80.5 | 80.5 | 80.5 |
| KP123 | ST512 | *phoE* | phoE_1 | 81 | 81 | 80.5 | 80.8 |
| KP123 | ST512 | *rpoB* | rpoB_1 | 83.5 | 83.5 | 83.5 | 83.5 |
| KP123 | ST512 | *tonB* | tonB_1 | 86.5 | 86.5 | 86.5 | 86.5 |
| KP123 | ST512 | *tonB* | tonB_2 | 88.5 | 88.5 | 88.5 | 88.5 |
| KP123 | ST512 | *gapA* | gapA_1 | 84.5 | 84.5 | 84.5 | 84.5 |
| KP123 | ST512 | *pgi* | pgi_1 | 86 | 86 | 86 | 86 |
| KP123 | ST512 | *wzi* | wzi_3 | 84 | 84 | 84 | 84 |
| KP123 | ST512 | *wzi* | wzi_4 | 83.5 | 83.5 | 83.5 | 83.5 |
| KP248 | ST423 | *infB* | infB_1 | 83.5 | 83.5 | 83.5 | 83.5 |
| KP248 | ST423 | *mdh* | mdh_1 | 80.5 | 80.5 | 80.5 | 80.5 |
| KP248 | ST423 | *phoE* | phoE_1 | 81 | 81 | 80.5 | 80.8 |
| KP248 | ST423 | *rpoB* | rpoB_1 | 83.5 | 83.5 | 83.5 | 83.5 |
| KP248 | ST423 | *tonB* | tonB_1 | 86.5 | 86.5 | 86.5 | 86.5 |
| KP248 | ST423 | *tonB* | tonB_2 | 89 | 89 | 89 | 89 |
| KP248 | ST423 | *gapA* | gapA_1 | 84.5 | 84.5 | 84.5 | 84.5 |
| KP248 | ST423 | *pgi* | pgi_1 | 86 | 86 | 86 | 86 |
| KP248 | ST423 | *wzi* | wzi_3 | 84 | 84 | 83.5 | 83.8 |
| KP248 | ST423 | *wzi* | wzi_4 | 83.5 | 83.5 | 83.5 | 83.5 |
| KP261 | ST258 | *infB* | infB_1 | 83.5 | 83.5 | 83.5 | 83.5 |
| KP261 | ST258 | *mdh* | mdh_1 | 80.5 | 80.5 | 80.5 | 80.5 |
| KP261 | ST258 | *phoE* | phoE_1 | 80.5 | 80.5 | 80.5 | 80.5 |
| KP261 | ST258 | *rpoB* | rpoB_1 | 83.5 | 83.5 | 83.5 | 83.5 |
| KP261 | ST258 | *tonB* | tonB_1 | 86.5 | 86.5 | 86.5 | 86.5 |
| KP261 | ST258 | *tonB* | tonB_2 | 88.5 | 88.5 | 88.5 | 88.5 |
| KP261 | ST258 | *gapA* | gapA_1 | 85 | 85 | 85 | 85 |

| Strain name | MLST profile | Gene name | Primer set name | Melting T 1 (°C) | Melting T 2 (°C) | Melting T 3 (°C) | Average melting T (°C) |
|---|---|---|---|---|---|---|---|
| KP261 | ST258 | *pgi* | pgi_1 | 86 | 86 | 86 | 86 |
| KP261 | ST258 | *wzi* | wzi_3 | 83.5 | 83.5 | 83.5 | 83.5 |
| KP261 | ST258 | *wzi* | wzi_4 | 84.5 | 84.5 | 84.5 | 84.5 |
| KP359 | ST149 | *infB* | infB_1 | 84 | 84 | 84 | 84 |
| KP359 | ST149 | *mdh* | mdh_1 | 80 | 80 | 80 | 80 |
| KP359 | ST149 | *phoE* | phoE_1 | 81 | 81 | 80.5 | 80.8 |
| KP359 | ST149 | *rpoB* | rpoB_1 | 84 | 84 | 84 | 84 |
| KP359 | ST149 | *tonB* | tonB_1 | 86.5 | 86.5 | 86.5 | 86.5 |
| KP359 | ST149 | *tonB* | tonB_2 | 89 | 89 | 89 | 89 |
| KP359 | ST149 | *gapA* | gapA_1 | 85 | 85 | 85 | 85 |
| KP359 | ST149 | *pgi* | pgi_1 | 86 | 86 | 86 | 86 |
| KP359 | ST149 | *wzi* | wzi_3 | 84 | 84 | 84 | 84 |
| KP359 | ST149 | *wzi* | wzi_4 | 84 | 84 | 84 | 84 |
| KP366 | ST11 | *infB* | infB_1 | 83.5 | 83.5 | 83 | 83.3 |
| KP366 | ST11 | *mdh* | mdh_1 | 80.5 | 80.5 | 80.5 | 80.5 |
| KP366 | ST11 | *phoE* | phoE_1 | 81 | 80.5 | 80.5 | 80.7 |
| KP366 | ST11 | *rpoB* | rpoB_1 | 83.5 | 83.5 | 83.5 | 83.5 |
| KP366 | ST11 | *tonB* | tonB_1 | 87 | 87 | 87 | 87 |
| KP366 | ST11 | *tonB* | tonB_2 | 88.5 | 88.5 | 88.5 | 88.5 |
| KP366 | ST11 | *pgi* | pgi_1 | 86 | 86 | 86 | 86 |
| KP366 | ST11 | *gapA* | gapA_1 | 85 | 85 | 85 | 85 |
| KP366 | ST11 | *wzi* | wzi_3 | 85 | 85 | 85 | 85 |
| KP366 | ST11 | *wzi* | wzi_4 | 84 | 84 | 84 | 84 |
| KP469 | ST395 | *infB* | infB_1 | 84 | 84 | 84 | 84 |
| KP469 | ST395 | *mdh* | mdh_1 | 80 | 80 | 80 | 80 |
| KP469 | ST395 | *phoE* | phoE_1 | 81 | 80.5 | 80.5 | 80.7 |
| KP469 | ST395 | *rpoB* | rpoB_1 | 83.5 | 83.5 | 83.5 | 83.5 |
| KP469 | ST395 | *tonB* | tonB_1 | 87 | 87 | 87 | 87 |
| KP469 | ST395 | *tonB* | tonB_2 | 89 | 89 | 89 | 89 |
| KP469 | ST395 | *gapA* | gapA_1 | 85 | 85 | 85 | 85 |
| KP469 | ST395 | *pgi* | pgi_1 | 85.5 | 85.5 | 85.5 | 85.5 |
| KP469 | ST395 | *wzi* | wzi_3 | 85 | 85 | 85 | 85 |
| KP469 | ST395 | *wzi* | wzi_4 | 83.5 | 83.5 | 83.5 | 83.5 |
| KP473 | ST101 | *infB* | infB_1 | 84 | 84 | 84 | 84 |
| KP473 | ST101 | *mdh* | mdh_1 | 80.5 | 80.5 | 80.5 | 80.5 |
| KP473 | ST101 | *phoE* | phoE_1 | 80.5 | 80.5 | 80.5 | 80.5 |
| KP473 | ST101 | *rpoB* | rpoB_1 | 83.5 | 83.5 | 83.5 | 83.5 |
| KP473 | ST101 | *tonB* | tonB_1 | 86.5 | 86.5 | 86.5 | 86.5 |
| KP473 | ST101 | *tonB* | tonB_2 | 89 | 89 | 88.5 | 88.8 |
| KP473 | ST101 | *pgi* | pgi_1 | 86 | 86 | 86 | 86 |
| KP473 | ST101 | *gapA* | gapA_1 | 85 | 85 | 85 | 85 |
| KP473 | ST101 | *wzi* | wzi_3 | 85 | 85 | 85 | 85 |
| KP473 | ST101 | *wzi* | wzi_4 | 84 | 84 | 83.5 | 83.8 |
| KP485 | ST15 | *infB* | infB_1 | 84 | 84 | 84 | 84 |
| KP485 | ST15 | *mdh* | mdh_1 | 80.5 | 80.5 | 80.5 | 80.5 |
| KP485 | ST15 | *phoE* | phoE_1 | 81 | 80.5 | 80.5 | 80.7 |

| Strain name | MLST profile | Gene name | Primer set name | Melting T 1 (°C) | Melting T 2 (°C) | Melting T 3 (°C) | Average melting T (°C) |
|---|---|---|---|---|---|---|---|
| KP485 | ST15 | *rpoB* | rpoB_1 | 83.5 | 83.5 | 83.5 | 83.5 |
| KP485 | ST15 | *tonB* | tonB_1 | 86.5 | 86.5 | 86.5 | 86.5 |
| KP485 | ST15 | *tonB* | tonB_2 | 89 | 89 | 89 | 89 |
| KP485 | ST15 | *gapA* | gapA_1 | 85 | 85 | 85 | 85 |
| KP485 | ST15 | *pgi* | pgi_1 | 86 | 86 | 86 | 86 |
| KP485 | ST15 | *wzi* | wzi_3 | 82.5 | 82.5 | 82 | 82.3 |
| KP485 | ST15 | *wzi* | wzi_4 | 84 | 84 | 84 | 84 |

**Table S2.** P-values of the Wilcoxon test (with Holm post-hoc correction) comparison of the average temperature distance matrices among the ten primer sets used in this work.

| | infB | mdh | phoE | rpoB | tonB_1 | tonB_2 | gapA | pgi | wzi_3 |
|---|---|---|---|---|---|---|---|---|---|
| **mdh** | 1.00000 (n.s) | | | | | | | | |
| **phoE** | 0.04820 (*) | 1.00000 (n.s) | | | | | | | |
| **rpoB** | 2.3e08 (****) | 0.00079 (***) | 0.00097 (***) | | | | | | |
| **tonB_1** | 1.00000 (n.s) | 1.00000 (n.s) | 1.00000 (n.s) | 0.00099 (***) | | | | | |
| **tonB_2** | 1.00000 (n.s) | 1.00000 (n.s) | 1.00000 (n.s) | 0.00050 (***) | 1.00000 (n.s) | | | | |
| **gapA** | 6.9e06 (****) | 0.02344 (*) | 0.07846 (n.s) | 1.00000 (n.s) | 0.03806 (*) | 0.02619 (*) | | | |
| **pgi** | 9.0e16 (****) | 9.9e09 (****) | 3.6e11 (****) | 0.04058 (*) | 3.9e09 (****) | 4.8e10 (****) | 0.00099 (***) | | |
| **wzi_3** | 0.00010 (***) | 0.00013 (***) | 3.6e06 (****) | 1.7e11 (****) | 0.00204 (**) | 0.00318 (**) | 2.8e10 (****) | <2e16 (****) | |
| **wzi_4** | 0.07276 (n.s) | 0.02344 (*) | 1.4e06 (****) | 4.8e13 (****) | 0.01799 (*) | 0.01594 (*) | 2.3e10 (****) | <2e16 (****) | 0.11752 (n.s) |

**Table S3.** Features and references of the *Klebsiella pneumoniae* strains used in this work.

| Strain name | Collection | MLST profile | wzi allele | Reference | Sequence Read Archive |
|---|---|---|---|---|---|
| 15PV | Background | ST976 | wzi_415 | Gaiarsa et al. 2015 | ERS480606 |
| 17PV | Background | ST307 | wzi_173 | Gaiarsa et al. 2015 | ERS480608 |
| 21PV | Background | ST240 | wzi_321 | Gaiarsa et al. 2015 | ERS480661 |
| 49BG | Background | ST147 | wzi_56 | Gaiarsa et al. 2015 | ERS480637 |
| 51BG | Background | ST1626 | wzi_417 | Gaiarsa et al. 2015 | ERS480639 |
| 52BG | Background | ST10 | wzi_95 | Gaiarsa et al. 2015 | ERS480640 |
| 53BG | Background | ST321 | wzi_131 | Gaiarsa et al. 2015 | ERS480641 |
| 55BG | Background | ST466 | wzi_37 | Gaiarsa et al. 2015 | ERS480643 |
| 63BG | Background | ST1627 | wzi_418 | Gaiarsa et al. 2015 | ERS480649 |
| BG-Kpn-20-18 | Outbreak | ST512 | wzi_154 | This work | Pending |
| BG-Kpn-21-18 | Outbreak | ST512 | wzi_154 | This work | Pending |
| BG-Kpn-22-18 | Outbreak | ST307 | wzi_173 | This work | Pending |
| BG-Kpn-23-18 | Outbreak | ST512 | wzi_154 | This work | Pending |

| Strain name | Collection | MLST profile | wzi allele | Reference | Sequence Read Archive |
|---|---|---|---|---|---|
| BG-Kpn-24-18 | Outbreak | ST512 | wzi_154 | This work | Pending |
| BG-Kpn-25-18 | Outbreak | ST512 | wzi_154 | This work | Pending |
| BG-Kpn-26-18 | Outbreak | ST512 | wzi_154 | This work | Pending |
| BG-Kpn-27-18 | Outbreak | ST512 | wzi_154 | This work | Pending |
| BG-Kpn-28-18 | Outbreak | ST512 | wzi_154 | This work | Pending |
| BG-Kpn-29-18 | Outbreak | ST512 | wzi_154 | This work | Pending |
| BG-Kpn-30-18 | Outbreak | ST512 | wzi_154 | This work | Pending |
| KP123 | Background | ST512 | wzi_154 | This work | Pending |
| KP248 | Background | ST423 | wzi_8 | This work | Pending |
| KP261 | Background | ST258 | wzi_29 | This work | Pending |
| KP359 | Background | ST149 | wzi_62 | This work | Pending |
| KP366 | Background | ST11 | wzi_75 | This work | Pending |
| KP469 | Background | ST395 | wzi_2 | This work | Pending |
| KP473 | Background | ST101 | wzi_137 | This work | Pending |
| KP485 | Background | ST15 | wzi_89 | This work | Pending |

**Table S4.** Clinical information of the outbreak strains.

| Strain name | Isolation date (yyyy/mm/dd) | Ward | Material |
|---|---|---|---|
| BG-Kpn-20-18 | 2018/04/10 | Infectious Diseases | Rectal swab |
| BG-Kpn-21-18 | 2018/04/10 | Infectious Diseases | Rectal swab |
| BG-Kpn-22-18 | 2018/04/11 | Oncology | Rectal swab |
| BG-Kpn-23-18 | 2018/04/15 | Oncology | Rectal swab |
| BG-Kpn-24-18 | 2018/04/16 | Infectious Diseases | Pus |
| BG-Kpn-25-18 | 2018/04/17 | Oncology | Rectal swab |
| BG-Kpn-26-18 | 2018/04/20 | Cardio-Surgical Intensive Care unit | Blood |
| BG-Kpn-27-18 | 2018/04/24 | Infectious Diseases | Rectal swab |
| BG-Kpn-28-18 | 2018/04/24 | Infectious Diseases | Urine |
| BG-Kpn-29-18 | 2018/04/24 | Oncology | Rectal swab |
| BG-Kpn-30-18 | 2018/04/26 | Cardio-Surgical Intensive Care unit | Rectal swab |

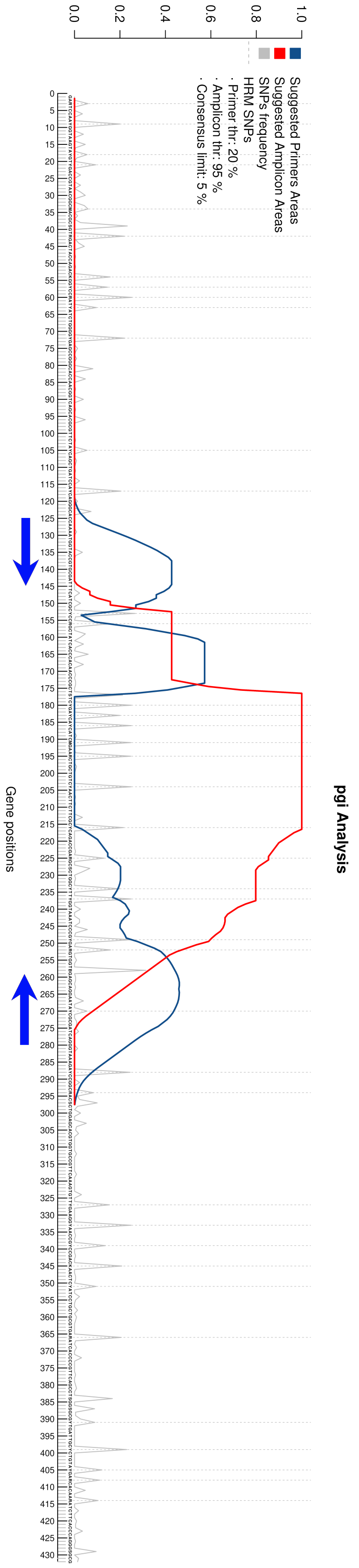


**Figure S1.** The output of EasyPrimer relative to the gene *pgi* (218 alleles on https://bigsdb.pasteur.fr). The consensus sequence calculated from the gene alignment is reported on the x axis. Residues under higher values of the blue curve are highly conserved and thus suitable for primer design, conversely the red curve increases over the highly variable regions suggested to be amplified. The grey peaks represent all the Single Nucleotide Polymorphisms (SNPs) with their own frequency. The dotted lines are used to highlight the "HRM-detectable" SNPs, so the one that cause a change in in GC content. The user can decide to consider only the "HRM-detectable" or all the SNPs for the analysis, thus highlighting the gene regions suitable for primer design for HRM and pan-PCR experiments respectively. The blue arrows were subsequently added manually to show the positions of the primer sets designed on *pgi* in this work.

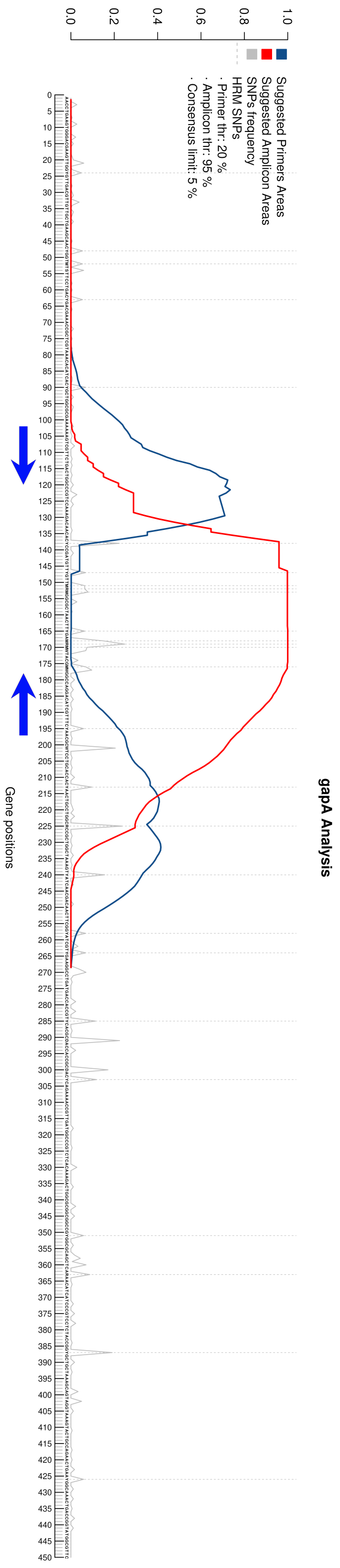


**Figure S2.** The output of EasyPrimer relative to the gene *gapA* (183 alleles on https://bigsdb.pasteur.fr). The consensus sequence calculated from the gene alignment is reported on the x axis. Residues under higher values of the blue curve are highly conserved and thus suitable for primer design, conversely the red curve increases over the highly variable regions suggested to be amplified. The grey peaks represent all the Single Nucleotide Polymorphisms (SNPs) with their own frequency. The dotted lines are used to highlight the “HRM-detectable” SNPs, so the one that cause a change in in GC content. The user can decide to consider only the “HRM-detectable” or all the SNPs for the analysis, thus highlighting the gene regions suitable for primer design for HRM and pan-PCR experiments respectively. The blue arrows were subsequently added manually to show the positions of the primer sets designed on *gapA* in this work.

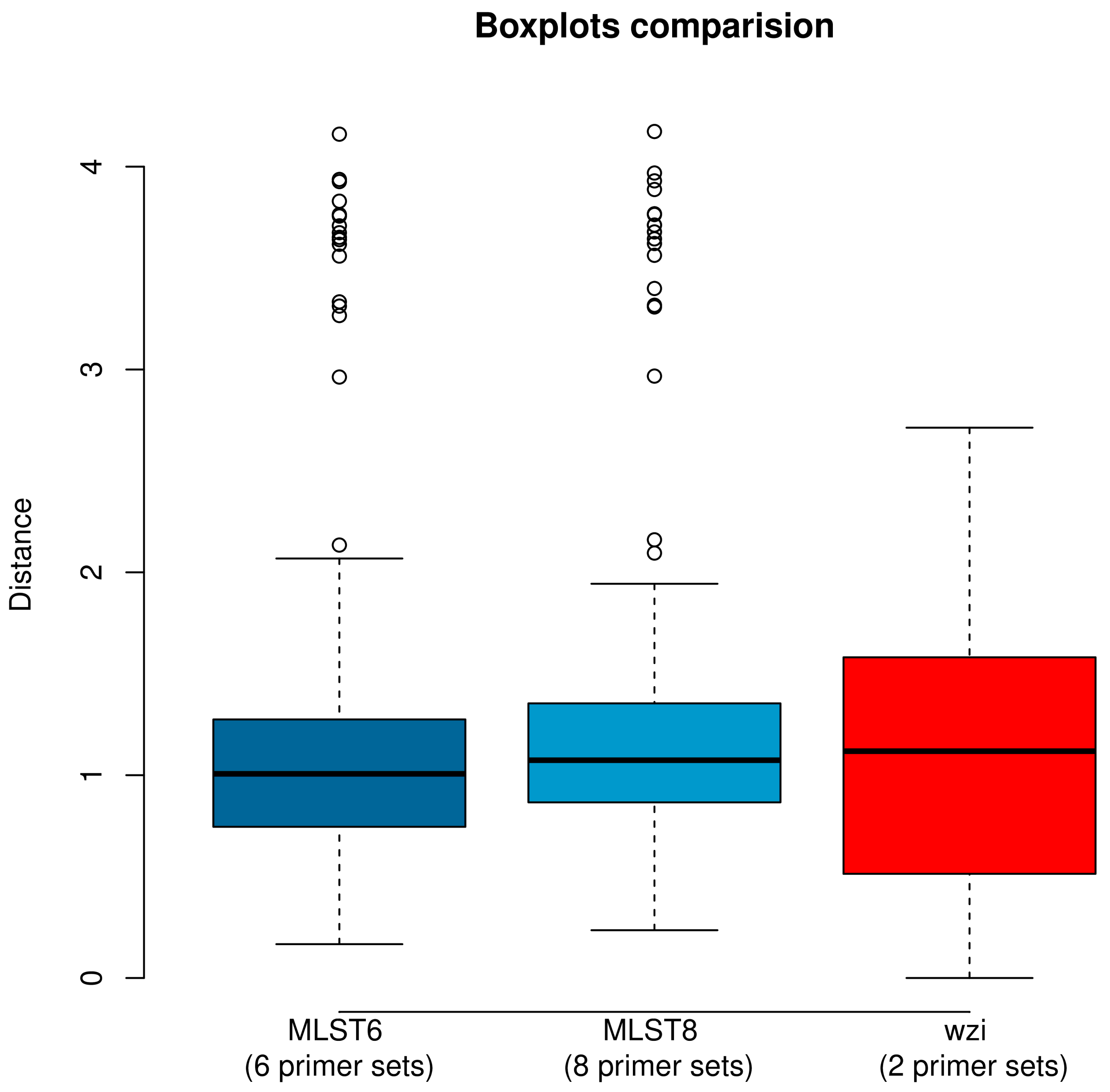


**Figure S3.** Distribution of the average melting temperature difference among the 17 *K. pneumoniae* strains for the three primer schemes. Boxes are the 25th and 75th quartiles divided by the medians, whiskers are 1.5x the interquartile ranges and dots are outliers.

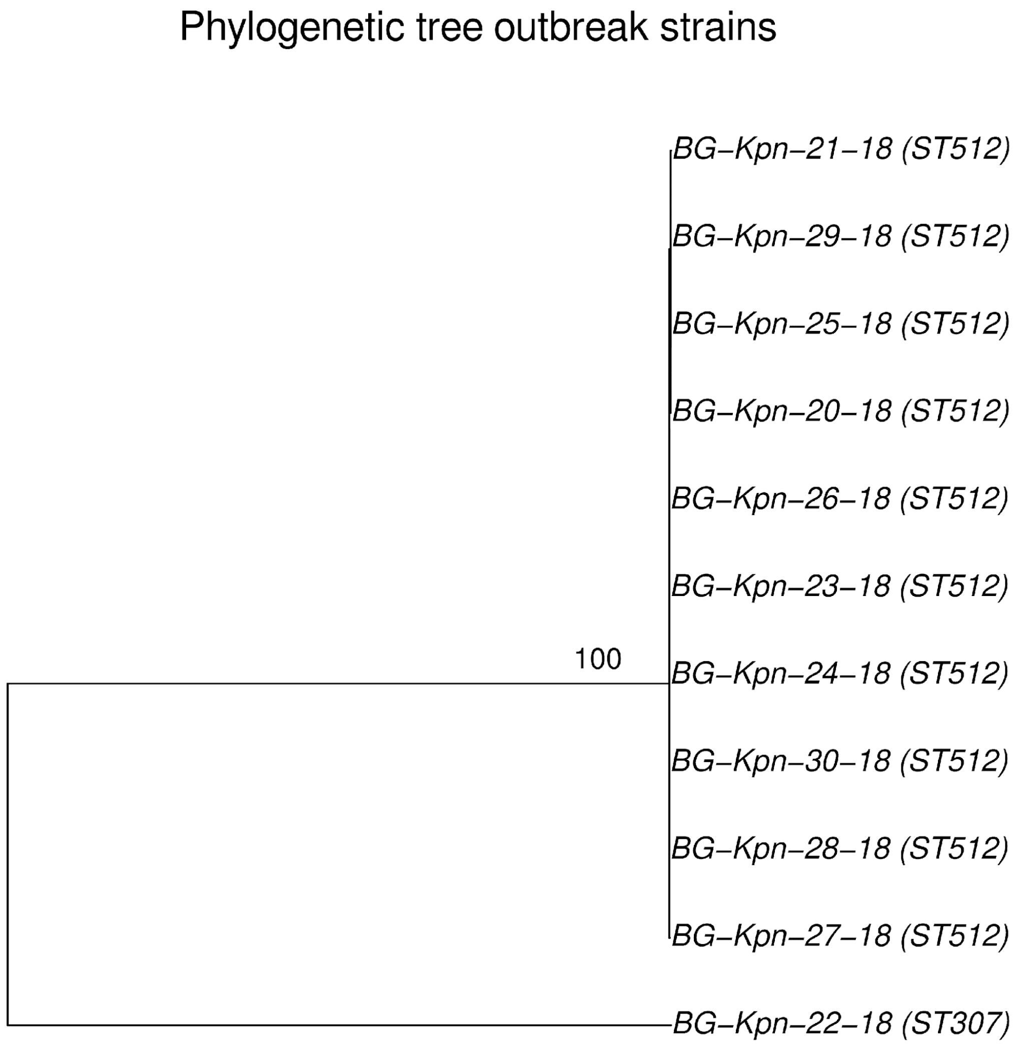


**Figure S4.** Maximum Likelihood phylogenetic tree of the 11 outbreak strains. The tree was obtained from an alignment of 66 core-SNPs.

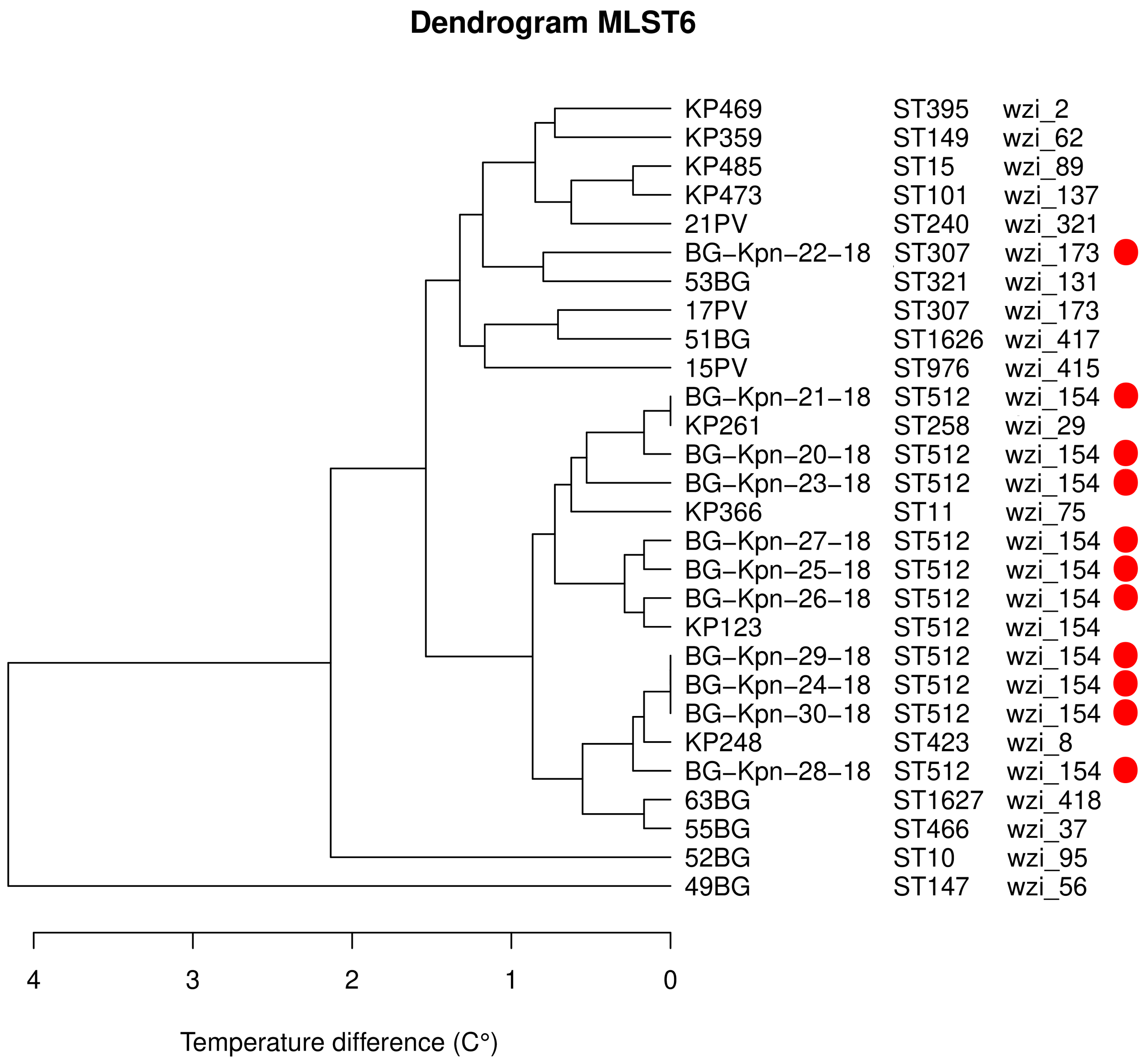


**Figure S6.** Dendrogram of the hierarchical clustering analysis on the average temperature distance matrix obtained using the MLST6 scheme. The 17 "background" strains belonging to 17 different MLSTs are written in black, while the 11 strains isolated during the nosocomial outbreak are highlighted by a red dot.

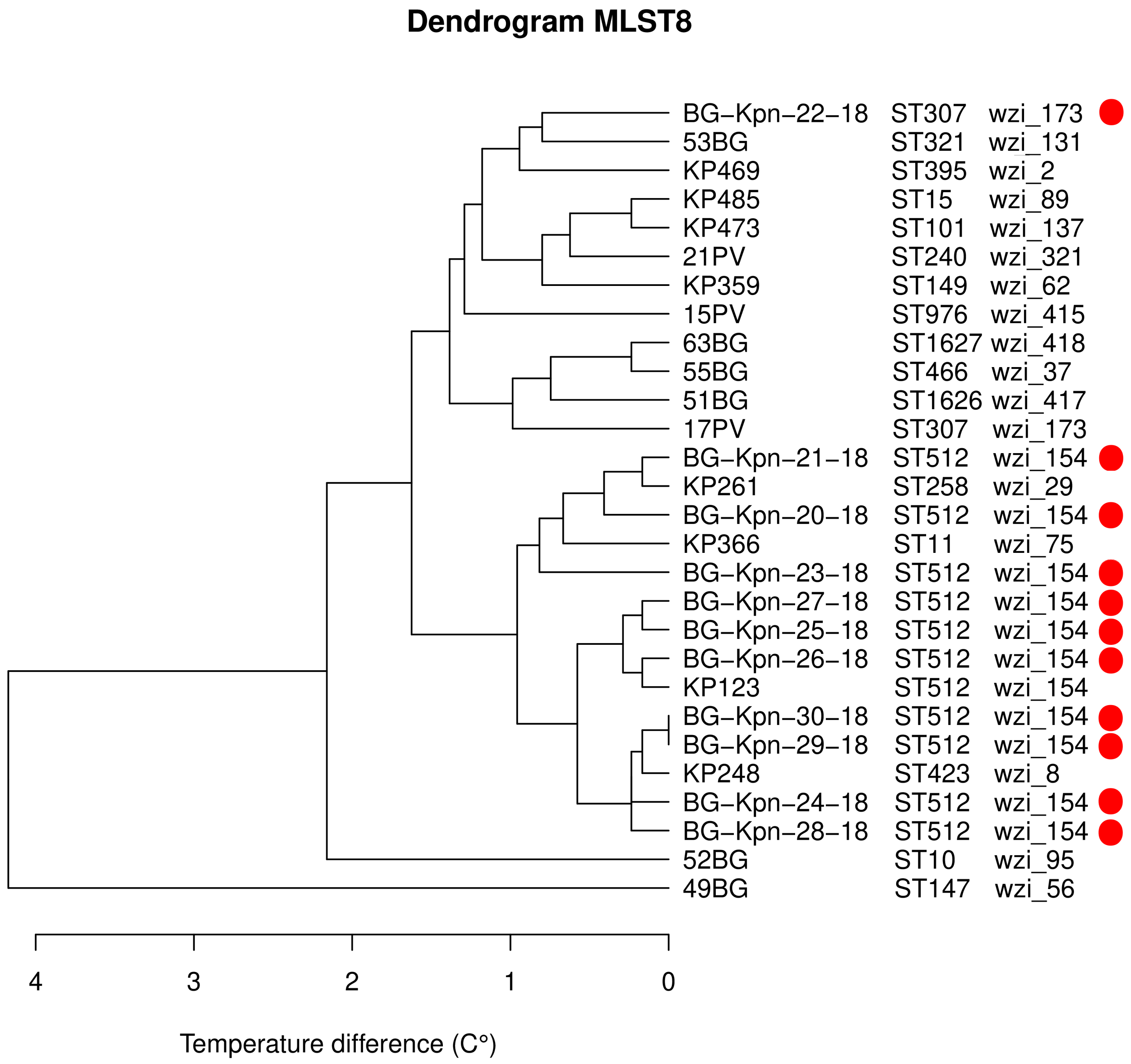


**Figure S7**. Dendrogram of the hierarchical clustering analysis on the average temperature distance matrix obtained using the MLST8 scheme. The 17 "background" strains belonging to 17 different MLSTs are written in black, while the 11 strains isolated during the nosocomial outbreak are highlighted by a red dot.

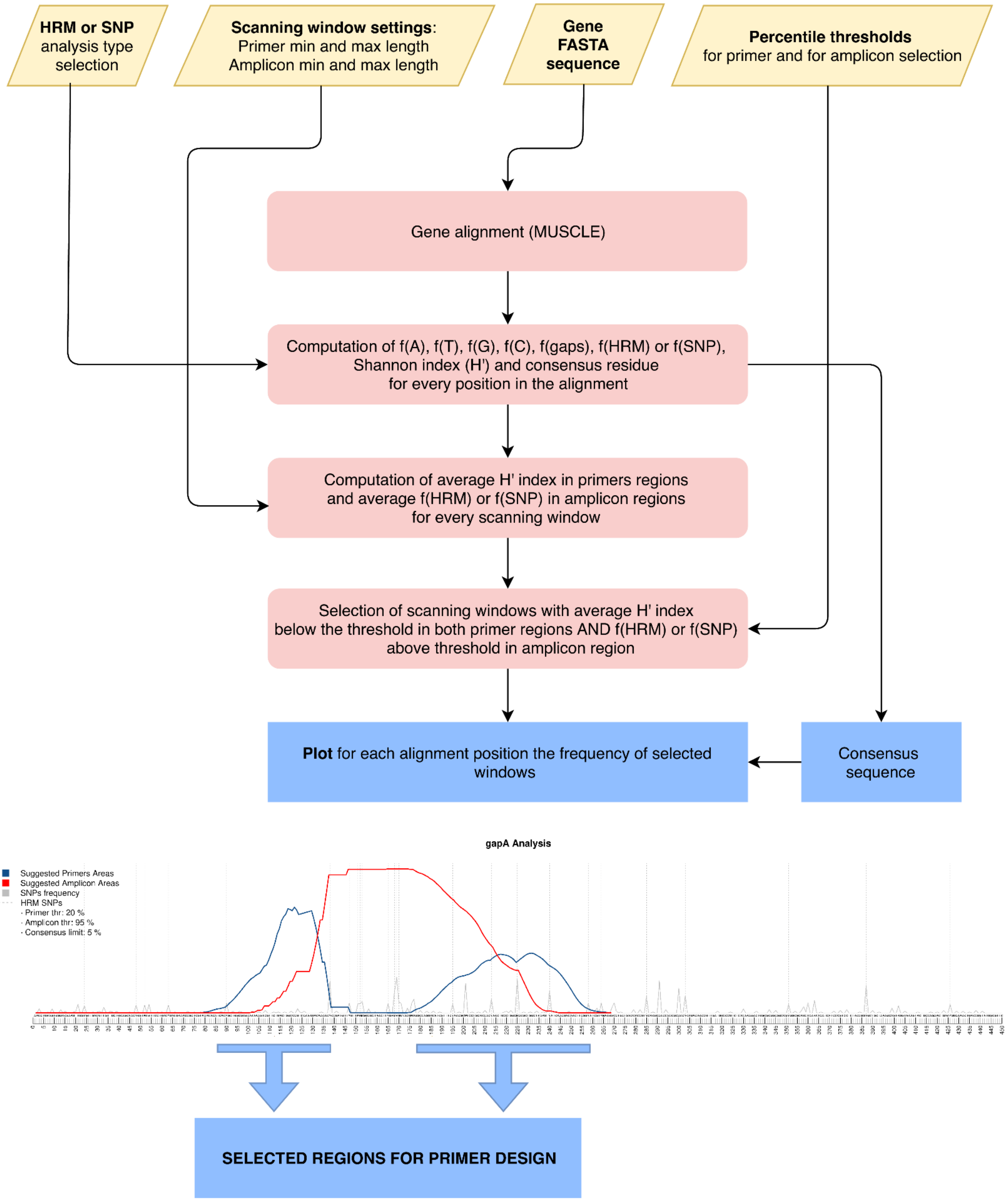


**Figure S7.** The diagram summarizes the main steps of the algorithm of EasyPrimer and shows an example of output.
